# Characteristics of early career health researchers and experiences of burnout during the COVID-19 pandemic in Canada

**DOI:** 10.1101/2024.11.13.622718

**Authors:** Sarah Hewko, Kaarina Kowalec, Laura Anderson, Erin E. Mulvihill, Maria J. Aristizabal, Annie V. Ciernia, Santokh Dhillon, Antoine Dufour, Gareth E. Lim, Maxime Rousseaux, Ayesha Saleem, Lubna Daraz, Grace Lam, the Association of Canadian Early Career Health Researchers (ACECHR)

## Abstract

**Introduction:** The COVID-19 pandemic disrupted research globally. How it impacted Canadian early-career health researchers (ECHRs) remains unclear. We administered a survey to understand the composition of ECHRs in Canada, their job experiences, and experiences of burnout during the COVID-19 pandemic.

**Methods:** A cross-sectional survey was conducted in May 2023 of Canadian ECHRs defined as within 7 years of their first independent research position. Quantitative analyses included a description of respondents by research pillar, socio-demographic and workplace characteristics, and the prevalence of burnout, disengagement or exhaustion. Sample characteristics were compared to national data on ECHRs from a Canadian funding agency. Thematic analysis of free-text responses was also conducted.

**Results:** A total of 225 respondents met the eligibility criteria. Most respondents were assistant professors and characteristics of our sample were like the national data. The COVID-19 pandemic posed many challenges to student recruitment, and emotional support of students, with over half of the respondents reporting a moderate to significant decline in mental health compared to pre-pandemic. A significant proportion of respondents were experiencing high burnout (62%, 95%CI:56-67%), exhaustion (64%, 95%CI: 57-70%) or disengagement (91%, 95%CI: 87-95%). Thematic analysis identified three themes: ongoing benefits/problems preceding the pandemic, unintended outcomes of strategies to manage/prevent/contain COVID-19, and reasons to stay in their current position.

**Conclusions:** Our survey revealed that Canadian ECHRs reported many diverse challenges during the COVID-19 pandemic and high burnout, putting the sustainability of this workforce at risk. Improved systems are needed to understand the long-term impacts and support the future of the Canadian health research ecosystem.

## Introduction

Health research is conducted by a diverse group of scientists, including undergraduate and graduate students, postdoctoral researchers, technical staff, clinicians, and faculty. “Early career health researcher” (ECHR) is a term used broadly to represent health researchers at the earliest stage of their academic research career.^1^ In the broader literature, the term often includes individuals within a specific number of years (typically 7) following completion of a PhD, MD or other terminal degree, meaning it can group together postdoctoral researchers and faculty.^1,2^ Given that postdoctoral researchers and faculty have diverse needs and face distinct career challenges, depending upon their work and geography, we sought to make a distinction between faculty and postdocs by specifically examining the challenges faced by faculty ECHRs (referred to as ECHR hereafter) in Canada. There is limited data on the demographics of Canadian ECHRs, information that is greatly needed to understand the challenges faced by this important group of health researchers, design interventions to support and retain ECHRs as they establish independent research careers and, ultimately, strengthen the Canadian health research ecosystem.

The COVID-19 pandemic disrupted research around the world and its lasting impacts are still not fully understood. The pandemic delayed research timelines, at least in part, by disrupting personnel recruitment and limiting access to laboratory space. It also forced researchers to deal with a variety of unexpected challenges including, among others, increased caregiving responsibilities in the home.^3–6^ In the USA, a survey of 284 science, technology, engineering, mathematics, and medicine faculty members revealed significant pandemic impacts, including fewer article submissions by women before and during the pandemic, an effect that is likely to have long-term consequences to career progression and that was more pronounced among faculty with young children.^2^ Focusing on ECHRs, Harrop et al. administered a survey to faculty, research scientists, and postdoctoral fellows working in the area of autism spectrum disorder; they reported similar effects, including negative impacts on research productivity, training opportunities, and mental health.^3^ Focusing on the latter, 65% of respondents reported at least one symptom of burnout, 17% reported persistent burnout, and 17% reported complete burnout. Importantly, this was a threefold increase in burnout compared to pre-pandemic levels, where almost 70% of respondents reported some stress, but no symptoms of burnout. In Canada, a survey of public health professionals also showed high rates of burnout during the pandemic. In this study, burnout, defined as “emotional exhaustion, depersonalization and diminished sense of achievement from the chronic exposure to stressors in the workplace”, was reported in 80% of respondents.^7^ Overall, the COVID-19 pandemic has significantly intensified burnout among early-career researchers and academics, magnifying pre-existing stressors and introducing new challenges that may disproportionally affect ECHRs.^3–6^

To understand the experience of Canadian ECHRs, the Association for Canadian Early Career Health Researchers (ACECHR) conducted a national survey. By collecting information about Canadian ECHR demographics, job characteristics and experiences, levels of burnout, and effects of the COVID-19 pandemic, we provide a baseline of information that can be used to inform, develop, implement, and evaluate future policies aimed at supporting ECHRs in Canada and beyond.

## Materials & Methods

### Data sources

#### Survey

The ACECHR National Steering Committee, made up of 11 ECHRs from across Canada, developed a 50-question survey (***Supplement***) to gain an understanding of ECHRs’ needs in Canada, and how their work-lives were impacted by the COVID-19 pandemic. The eligible respondents were currently independent health researchers at Canadian institutions and had been in such a role for no more than seven years. English-language fluency was necessary to complete the survey.

In the survey, respondents were asked to provide information on demographic characteristics, details of their research funding history and perceived impacts of COVID-19 on their work-lives. Also included in the survey was the Oldenburg Burnout Inventory (OLBI). The survey was hosted on LimeSurvey and could be completed anonymously in approximately 20 minutes. The only questions for which a response was mandatory were those required to establish eligibility for the study. The survey was open from April 28, 2023 - May 28, 2023. Respondents were recruited to participate through social media posts on multiple platforms. In addition to posts shared on the ACECHR Twitter/X account, an e-mail was sent from the Co-Chairs of the ACECHR National Committee (using the ACECHR e-mail) to encourage participation from current members of ACECHR (n=203). Individual members of the ACECHR National Committee shared recruitment posts on their personal social media accounts and facilitated sharing of survey information in targeted newsletters and communications from institutions and agencies with which they were associated. The protocol for the study was approved by the University of Prince Edward Island Research Ethics Board for compliance with Canadian guidelines for research involving human participants (File #6011967).

#### Canadian Institutes of Health Research (CIHR) Data

For comparison, we obtained ECHR funding data from CIHR. CIHR is one of the “tri-agency” funding agencies. Tri-agency is an umbrella term to describe the three funding agencies of the Canadian government which span health research, natural sciences and engineering, and social sciences and humanities. Twice annually CIHR offers Project Grant competitions, which are a significant potential source of funding for health researchers, including ECHRs, in Canada. The nominated principal applicant (NPA) (or principal investigator) must be affiliated with a Canadian postsecondary institution and/or an affiliated institution (e.g., hospitals or research institutes), or be an individual or organization affiliated with an Indigenous non-governmental organization within Canada that has a research and/or knowledge translation mandate. At our request in July 2023, CIHR provided summary-level data about applicants to the 2019-2022 Project Grant competitions, which included: the total number of unique ECHRs who applied as NPA, split by research pillar (CIHR organizes health research into four pillars – biomedical; clinical; health services; and social, cultural, environmental and population health); gender identity (gender-fluid, man, woman, two-spirit, nonbinary, queer, or prefer not to answer), and; self-identified disability (yes, no, or prefer not to answer). As per CIHR procedures, any cells with counts N<10 were suppressed to maintain privacy. In cases where data from a single group was N<10, values for those who responded ’I prefer not to answer’ were also suppressed.

### Quantitative Analysis

We described the characteristics of eligible survey respondents. For continuous variables, the median and standard deviation (SD) were reported, and for categorical variables the frequency and percent were reported. Descriptive statistics were reported overall and stratified by research pillar, institution type, and gender identity. For work location, we combined six of the ten provinces into two categories: Prairies (Manitoba, Saskatchewan) and Atlantic Canada (New Brunswick, Nova Scotia, Prince Edward Island, and Newfoundland/Labrador). The remaining provinces (Alberta British Columbia, Ontario, Québec) were examined independently. Summary-level data from CIHR were provided post-descriptive analysis and were organized by pillar, gender, and self-identified disability. Burnout was measured using the Oldenburg Burnout Inventory (OLBI), which is a validated 16-item measure of burnout that consists of two subscales (exhaustion and disengagement).^8^ The tool has been applied to study burnout in similar populations, including among health science faculty in the USA,^9^ academics at a tertiary institution in South Africa^10^ and a global sample of academic radiographers.^11^ Knapp et al. reported a Cronbach’s alpha coefficient of 0.83 for the OLBI (including both dimensions of burnout) in their sample. Each subscale consisted of 8 items - four were positively worded and four were negatively worded - with four-point Likert scale response options ranging from 1=totally disagree to 4=totally agree. Scores were calculated overall and for each subscale as the mean sum of the items with the negatively coded responses reverse coded. High disengagement was defined as ≥2.1 and high exhaustion was defined as ≥2.25.^12^ High burnout was defined as the presence of both high disengagement and high exhaustion. Unadjusted regression analyses tested the association between four candidate factors determined a priori (age, gender identity, years of independence, childcare duties) and three outcomes: high burnout, high disengagement, and high exhaustion. Unadjusted odds ratios (OR) and 95% confidence intervals (CI) were calculated from logistic regression.

Quantitative analyses were conducted using SAS (v9.4), R for Statistical Computing (v4.3.1), R-Studio (v2024.04.1) and GraphPad-Prism software (v10.1.2, San Diego, CA, USA). Missing data were not imputed. Statistical significance was indicated by p≤0.05.

### Qualitative Analysis

We conducted a thematic analysis of responses to the survey’s final, free text question which queried if there was anything else respondents wanted to share about their experience as an ECHR during the COVID-19 pandemic. All responses to the question (n=80) were independently reviewed and inductively coded in MAXQDA by two authors (SH and EEM) familiar with the survey and with the results of the quantitative descriptive analysis. Segments of text fitting with multiple codes were coded with as many codes as were relevant. Having completed independent coding, SH and EEM met to discuss all assigned codes. Together, based on consensus coding, they agreed on a thematic structure that best represented the qualitative findings. The relative importance of a theme or subtheme was determined based on the prevalence of related codes in the data. Codes that appeared fewer than three times in the data were eliminated in the final thematic structure.

## Results

### Demographics and job characteristics of Canadian ECHRs

Out of 264 complete responses, 225 respondents met all eligibility criteria - ≤7 years as an independent health researcher at a Canadian institution (Table 1). The median age of the respondents was 39.5 years, 62% identified as women (n=139, Figure 1A and B), and 64% reported having children, with the majority having 2 young children (51%, Table 2). Across research pillars (Table 1), institution types (Table S1) and gender (Table S2), characteristics for the survey respondents were broadly like the full sample of ECHR survey respondents.

**Figure 1:**
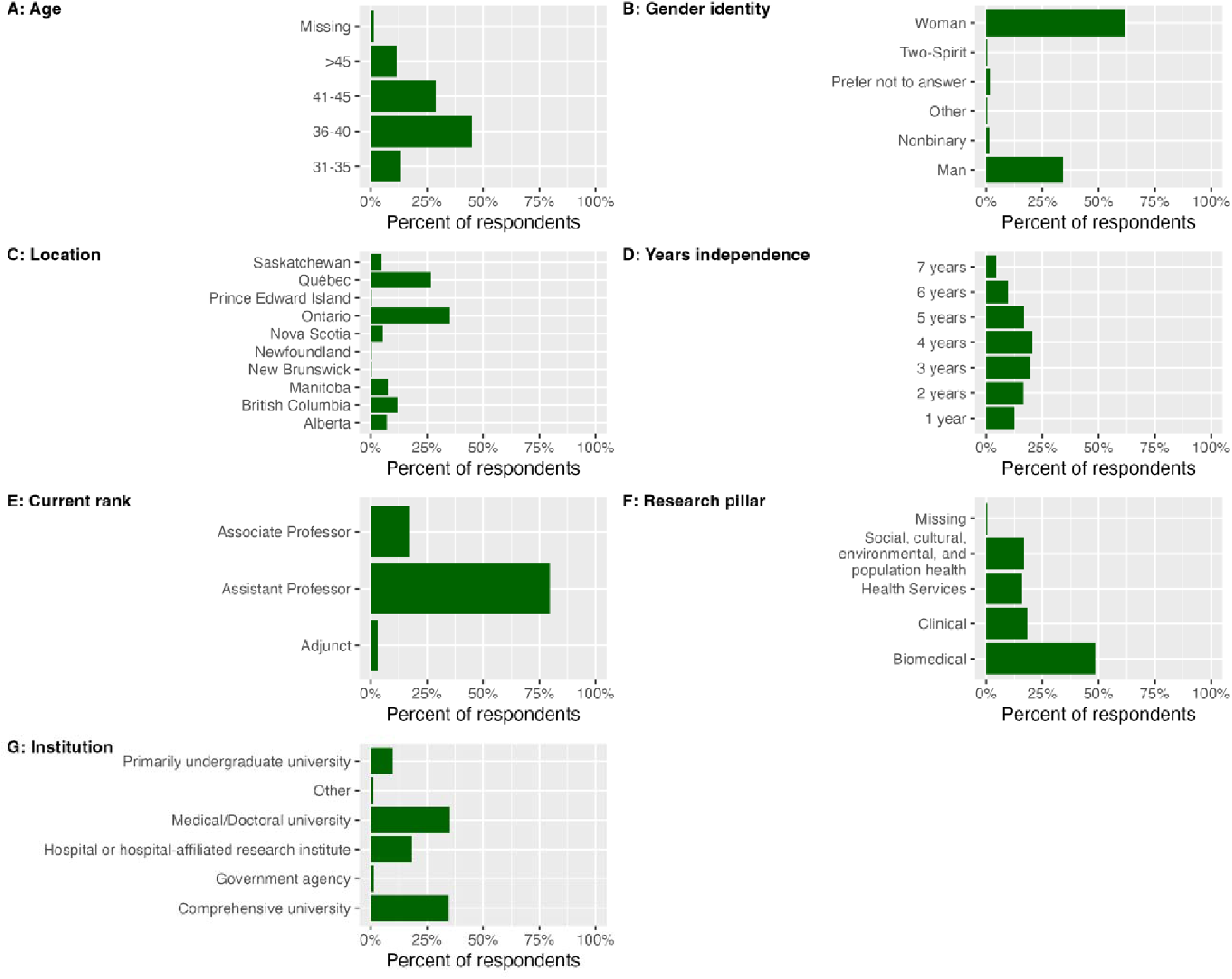
Demographic and professional characteristics of Canadian ECHR respondents. Bar graphs showing (A) age, (B) gender identity, (C) Canadian province where respondent works, (D) number of years as an independent investigator, (E) current academic rank, (F) research pillar that best describes the ECHR work and (G) type of academic institution at which respondents are employed.

**Table 1:** Job characteristics and demographics of Canadian early career health researchers, by research pillar.

|  | Entire sample,<br>N=225 | Research Pillar<sup>A</sup> |  |  |  |
| --- | --- | --- | --- | --- | --- |
|  |  | Biomedical,<br>n=109 | Clinical,<br>n=41 | Health services,<br>n=36 | Social, cultural,<br>environmental, and<br>population health, n=38 |
| **Age, years** | 39.5 (37, 43) | 40 (37, 42) | 38.5 (36, 44) | 39 (36, 44.2) | 39 (37, 44) |
| **Gender Identity** |  |  |  |  |  |
| Woman | 139 (62.0) | 54 (49.5) | 28 (68.3) | 25 (69.4) | 32 (84.0) |
| Man | 77 (34.0) | 50 (45.9) | 12 (29.3) | 9 (25.0) | 5 (13.0) |
| Mix of gender identities | 5 (2.0) | ¶ | ¶ | ¶ | ¶ |
| Prefer not to answer | ¶ | ¶ | ¶ | ¶ | ¶ |
| **Independent position, years** | 4 (2, 5) | 3 (2, 5) | 4 (3, 4) | 4 (2, 5) | 4 (2, 5) |
| **Province or territory<sup>B</sup>** |  |  |  |  |  |
| Alberta | 16 (7.1) | 10 (9.2) | ¶ | ¶ | ¶ |
| British Columbia | 27 (12.0) | 11 (10.1) | ¶ | 5 (13.9) | 8 (21.0) |
| Prairies | 28 (12.4) | 17 (15.7) | ¶ | ¶ | ¶ |
| Atlantic Canada | 15 (6.7) | 8 (7.3) | ¶ | ¶ | ¶ |
| Ontario | 79 (35.1) | 36 (33.0) | 15 (36.5) | 17 (47.2) | 11 (28.9) |
| Québec | 60 (26.6) | 27 (24.8) | 17 (41.4) | 6 (16.7) | 9 (23.6) |
| **Current rank** |  |  |  |  |  |
| Assistant Professor | 179 (79.5) | 90 (82.6) | 34 (82.9) | 27 (75) | 27 (71.1) |
| Associate Professor | 39 (17.3) | 15 (13.8) | 7 (17.1) | 8 (22.2) | 9 (23.7) |
| Adjunct | 7 (3.1) | ¶ | 0 | ¶ | ¶ |
| **Institute type<sup>C</sup>** |  |  |  |  |  |
| Comprehensive | 78 (34.6) | 46 (42.2) | 10 (24.4) | 9 (25) | 13 (34) |
| Medical/doctoral | 79 (35.1) | 38 (34.8) | 14 (34.2) | 15 (42) | 11 (29) |
| Hospital | 41 (18.2) | 9 (8.3) | 16 (39) | 7 (19) | 9 (24) |
| Primarily undergraduate | 22 (9.7) | 14 (12.8) | ¶ | ¶ | ¶ |
| **Job characteristics** |  |  |  |  |  |
| Teaching stream | 22 (9.7) | 6 (5.5) | 10 (24.4) | ¶ | ¶ |
| Clinical stream | 37 (16.4) | 5 (4.5) | 19 (46.3) | 8 (22.2) | ¶ |
| **Grant agencies applied to** |  |  |  |  |  |
| Tri-agency | 204 (90.6) | 100 (91.7) | 37 (90.2) | 30 (83.3) | 36 (94.7) |
| National agency | 111 (49.3) | 67 (61.5) | 12 (29.2) | 16 (44.4) | 15 (39.4) |
| Non-profit/charity | 145 (64.4) | 82 (75.2) | 25 (60.9) | 19 (52.8) | 18 (47.4) |
| Province/territory | 150 (66.6) | 74 (67.9) | 28 (68.3) | 21 (58.3) | 26 (68.4) |
| US agency | 62 (27.5) | 41 (37.6) | 11 (26.8) | ¶ | 6 (15.7) |
| International not US | 33 (14.6) | 18 (16.5) | 7 (17.1) | ¶ | ¶ |

Numbers are reported as n (%) or median (interquartile range), ¶ = Suppressed as N<5, A: 1 person did not report a research pillar. B: No respondents from a Canadian territory. C: Institute type of Government or Other: N<5.

**Table 2:**
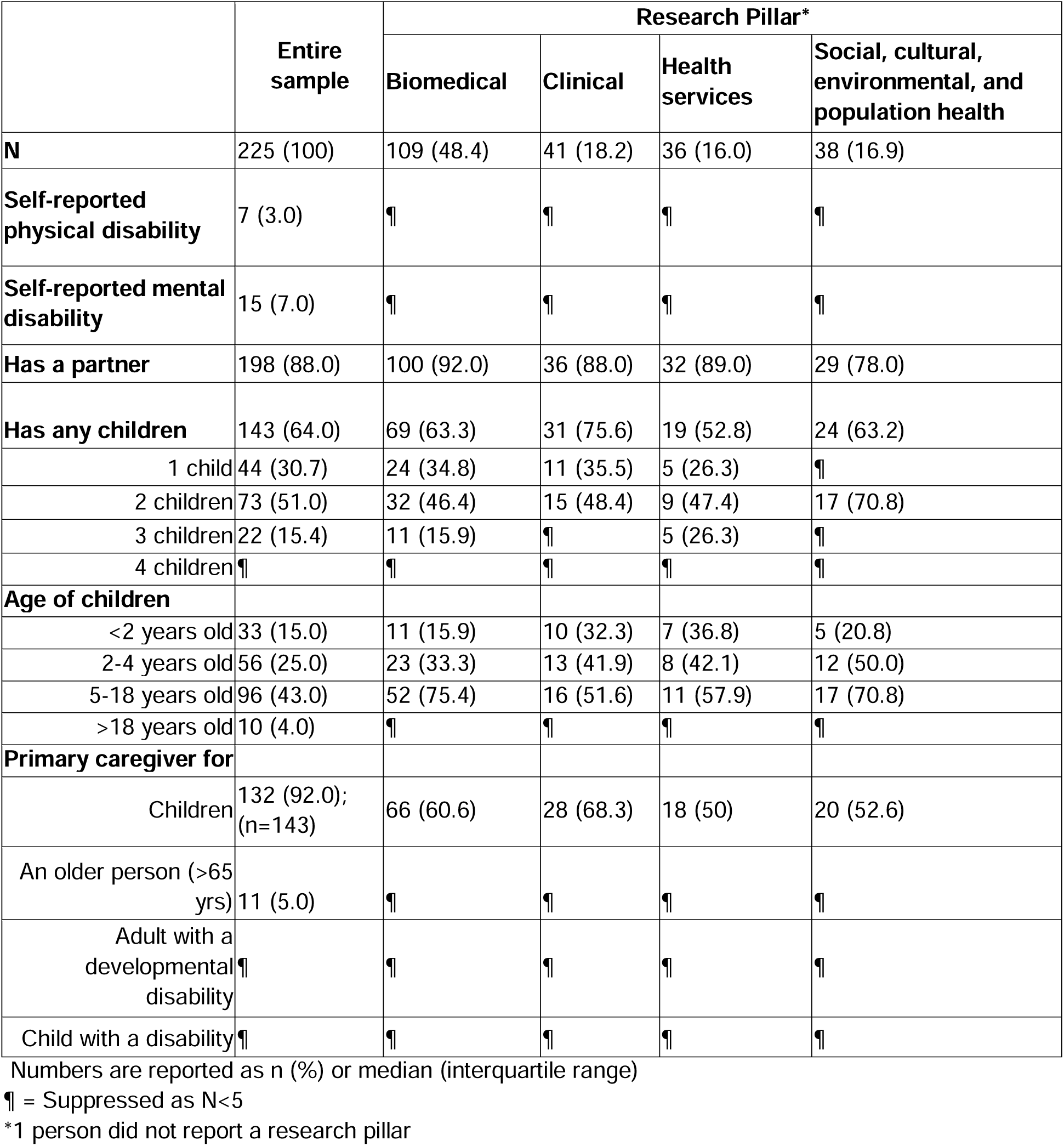
Additional characteristics of Canadian early career health researchers, by research pillar.

Importantly, the survey captured ECHRs during the first 7 years of their independent career and working across all Canadian provinces although most respondents worked in Ontario and Québec (>60%), the two most populated provinces in Canada. Overall, we found that most respondents primarily conducted research that falls into the biomedical research pillar category (48%, Figure 1F). Most respondents were employed at comprehensive (undergraduate and graduate) or medical/doctoral universities (70%) at the rank of assistant professor (80%, Figure 1). Workloads varied widely with research responsibilities showing the largest variation (25^th^ and 75^th^ percentiles of 40-75%), followed by clinical (30-50%), teaching (10-30%) and service roles (10-20%, Figure 2).

**Figure 2.**
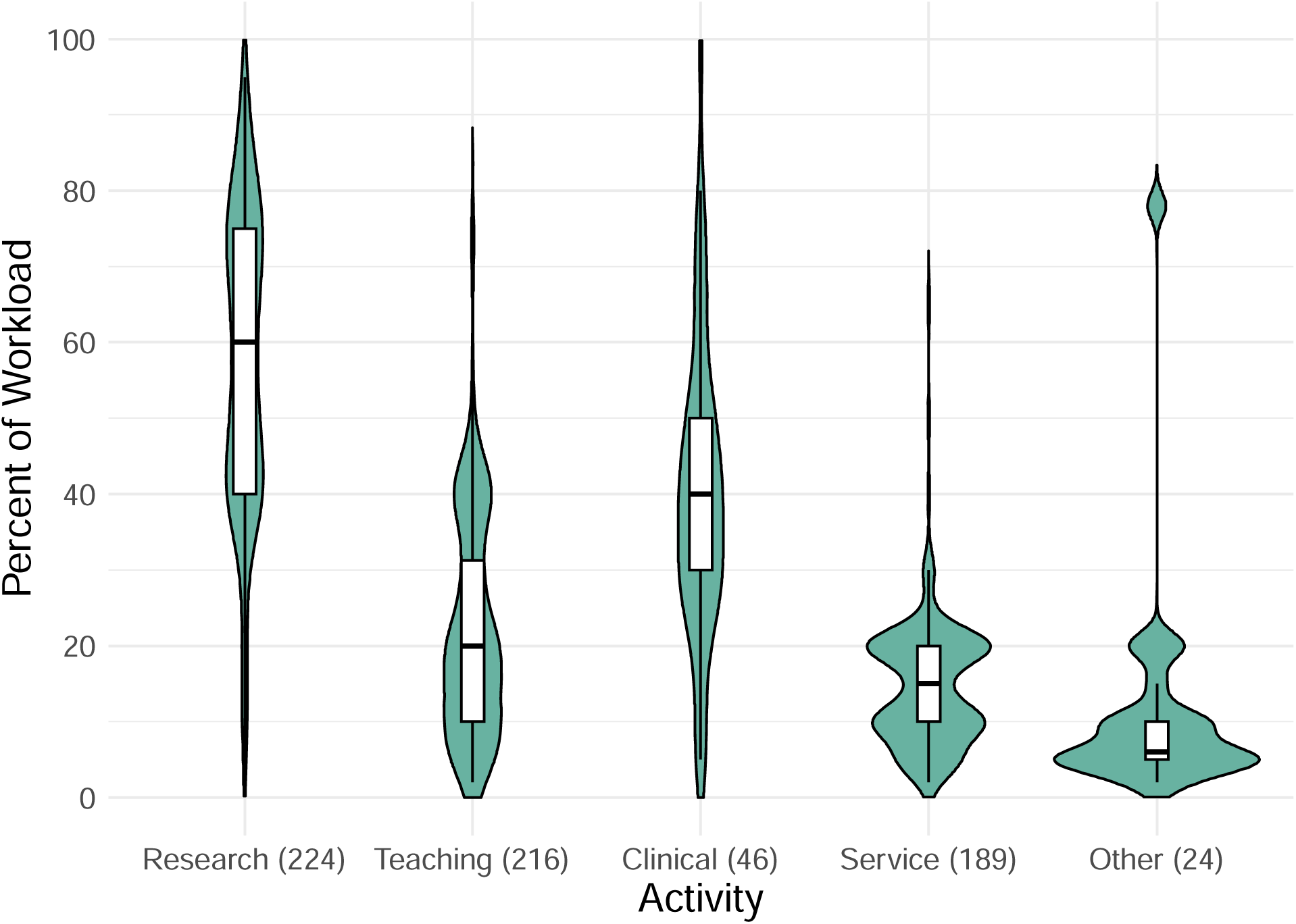
Workload distribution Canadian ECHR survey respondents. Violin plot showing the workload distribution of Canadian ECHRs. Boxplots show the median workload, with lower and upper edges showing the first (25th percentile) and third (75th percentile) quartiles, respectively.

### ECHR research funding

Among ECHRs, research funding was generally sought from national research agencies (91%), national co-agencies (49%), non-profits/charities (65%), and provinces or territorial research organizations (67%, Table 1). From 2019-2022, the mean number of applications as principal investigator or principal applicant to either tri-council or non-tri-council agencies was approximately one application annually, but the range was from zero applications to as many as 20 applications in a single year (Table 3). The average number of successful applications, which included applications as primary investigator, or in any other role (e.g., co-investigator, collaborator), was 1.2-1.5 applications annually. The top of the range for successful applications annually was 16 (Table 3).

**Table 3:** Number of grant applications submitted as principal investigator or principal applicant.

|  | Year |  |  |  |
| --- | --- | --- | --- | --- |
|  | 2019 (n=224) | 2020 (n=221) | 2021 (n=218) | 2022 (n=220) |
| Number of applications submitted to tri-council agencies (Canadian government funding) | 1.0 (1.4, 0-20) | 0.9 (1.3, 0-9) | 1.1 (1.2, 0-6) | 1.3 (1.5, 0-14) |
| Number of applications submitted to other agencies | 1.7 (2.1, 0-11) | 1.6 (2.1, 0-19) | 1.6 (2.0, 0-13) | 1.8 (2.0, 0-12) |
| Number of successful applications (overall) | 1.2 (1.6, 0-8) | 1.2 (1.4, 0-7) | 1.3 (1.5, 0-12) | 1.5 (1.5, 0-16) |

All values are reported as the mean number of applications (standard deviation, range).

To determine if our survey was representative of ECHRs that apply for Canadian national health research funding, we compared the demographics of our survey respondents to demographic information obtained from the national Canadian agency (CIHR) for ECHRs that applied to the national open competition (Project Grant) between 2019-2022 (Table 4). CIHR runs two Project Grant competitions per year, in fall and spring, which are examined separately here. On average, approximately 500 ECHRs applied as the Nominated Principal Applicant (PI) to either of the biannual CIHR Project Grant competitions, compared to 204 out of 225 (90.6%) respondents to our survey, who reported applying to one of the Canadian governmental (tri-agency) competitions (Table 1). Across research pillars (Table 1 and 3), institution types (Table S1) and gender (Table S2), characteristics for the survey respondents were like that of those ECHRs that had recently applied to CIHR for research funding. However, our survey included more ECHRs that identified as women (62% compared to 48% among ECHRs that applied for CIHR funding, Table 4) and that self-reported a disability (8% compared to 3% among ECHR that applied for CIHR funding, Table 4). For the latter, we do note an increase in the proportion with a self-reported disability in the 2022 Fall Project Grant competition (5.7%).

**Table 4:**
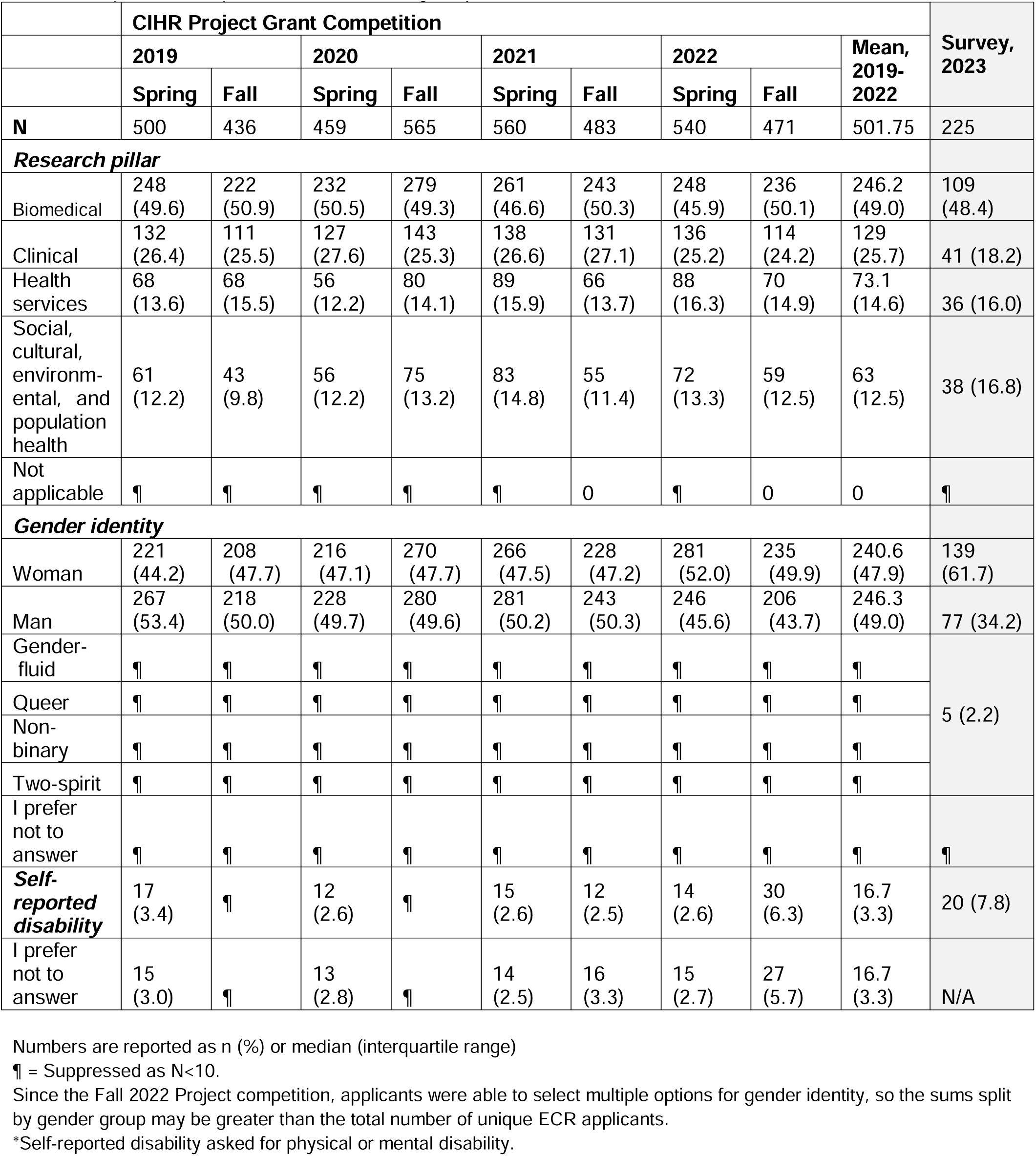
Demographics of Canadian early career health researcher applicants, by CIHR Project Grant Competition compared to current survey respondents.

We noted that several ECHRs reported having submitted fewer grant applications as principal investigator or co-investigator than expected. The main reasons for not applying to as many grants as expected included: inadequate resources (40%), feeling discouraged (29%), and not being ready (29%), responses that were similar in proportion across research pillar, gender identity, and institution type (Table S3).

### COVID-19 pandemic-related effects on Canadian ECHRs

Our survey also included questions aimed at understanding the impacts of the COVID-19 pandemic on the ECHR experience and to evaluate the effectiveness of policies that were implemented to mitigate its effects. More than half (51%) of the respondents indicated that their institution offered an extension to the tenure clock because of the COVID-19 pandemic (Table 5) with most (89.4%) indicating that the extension was for one year. Of those aware of an extension being offered, only 26% indicated they were likely to request this extension when it came time for their tenure application (Table 5). ECHRs research direction also changed because of the pandemic, with around half of the respondents reporting having pivoted at least partially to COVID-19 related research during this time. More than half of the respondents reported having moderately to significantly worse mental health at the time of survey compared to pre-pandemic levels (n=146, 64.8%).

**Table 5:** COVID-19 related descriptives by the entire sample and by research pillar.

|  | Total | Research Pillar |  |  |  |
| --- | --- | --- | --- | --- | --- |
|  |  | Biomedical | Clinical | Health services | Social, cultural, environmental, and population health |
| **N** | 225 | 109 (48.4) | 41 (18.2) | 36 (16) | 38 (16.9) |
| **COVID-19 time extension for tenure** |  |  |  |  |  |
| Yes | 114 (50.7) | 62 (56.8) | 16 (39.0) | 19 (52.8) | 17 (44.7) |
| No | 37 (16.4) | 19 (17.4) | 7 (17.1) | ¶ | 6 (15.8) |
| University does not have tenure track | 11 (4.8) | ¶ | 5 (12.2) | ¶ | ¶ |
| I don't know | 59 (26.2) | 24 (22.0) | 13 (31.7) | 8 (22.2) | 14 (36.8) |
| Prefer not to answer | ¶ | ¶ | 0 | ¶ | 0 |
| **COVID-19 length of extension (n=114)** |  |  |  |  |  |
| 1 year | 102 (89.4) | 58 (93.5) | 15 (93.8) | 15 (78.9) | 14 (82.4) |
| 2 years | 12 (10.5) | ¶ | ¶ | ¶ | ¶ |
| **COVID-19 tenure extension used (n=113)** |  |  |  |  |  |
| Definitely no (or didn't) | 50 (44.2) | 25 (40.3) | 5 (31.2) | 10 (52.6) | 10 (58.8) |
| Probably no | 19 (16.8) | 10 (16.1) | ¶ | ¶ | ¶ |
| Have not decided yet | 14 (12.4) | 8 (12.9) | ¶ | ¶ | ¶ |
| Probably yes | 8 (7.0) | 6 (9.7) | ¶ | 0 | 0 |
| Definitely yes | 22 (19.5) | 13 (21) | ¶ | 5 (26.3) | 0 |
| **Impact of citizenship status on ECHR status during COVID-19 (n=219)** |  |  |  |  |  |
| Impacted my experience very negatively | 16 (7.3) | 10 (9.2) | ¶ | ¶ | ¶ |
| Impacted my experience negatively | 24 (10.9) | 15 (13.8) | ¶ | ¶ | 5 (13.2) |
| Had no impact on my experience | 168 (76.7) | 80 (73.4) | 34 (82.9) | 27 (75) | 26 (68.4) |
| Impacted my experience positively | 8 (3.7) | ¶ | ¶ | ¶ | ¶ |
| Impacted my experience very positively | 3 (1.4) | 0 | 0 | ¶ | ¶ |
| **Mental health changes during COVID-19** |  |  |  |  |  |
| Significantly worsened | 36 (16.0) | 22 (20.2) | 5 (12.2) | ¶ | 7 (18.4) |
| Moderately worsened | 110 (48.8) | 49 (45) | 18 (43.9) | 21 (58.3) | 21 (55.3) |
| Has not changed | 67 (29.8) | 32 (29.4) | 16 (39) | 11 (30.6) | 8 (21.1) |
| Moderately improved | 10 (4.4) | 6 (5.5) | ¶ | ¶ | ¶ |
| Improved significantly | ¶ | 0 | 0 | ¶ | 0 |
| Prefer not to answer | ¶ | 0 | ¶ | 0 | 0 |
| **Pivoted to COVID-19 research** |  |  |  |  |  |
| No | 111 (49.3) | 76 (69.7) | 17 (41.5) | 10 (27.8) | 8 (21.1) |
| Partially | 103 (45.8) | 28 (25.7) | 24 (58.5) | 24 (58.5) | 26 (68.4) |
| Fully | 9 (4.0) | ¶ | 0 | ¶ | ¶ |

Numbers are reported as n (%) or median (interquartile range). ¶ = Suppressed as N<5.

Respondents who had been in an independent research position prior to the COVID-19 pandemic (n=116) indicated that recruiting students (67%), providing emotional support for students and trainees (66%), and teaching (54%) were more difficult now than pre-COVID (Figure 3a). Those who began their independent positions during the COVID-19 pandemic indicated that recruiting students (69%), communication with administration (47%), ordering supplies (48%), emotional support of students/trainees (46%) and accessing resources (44%) had proven to be more difficult than they had expected (Figure 3b).

**Figure 3:**
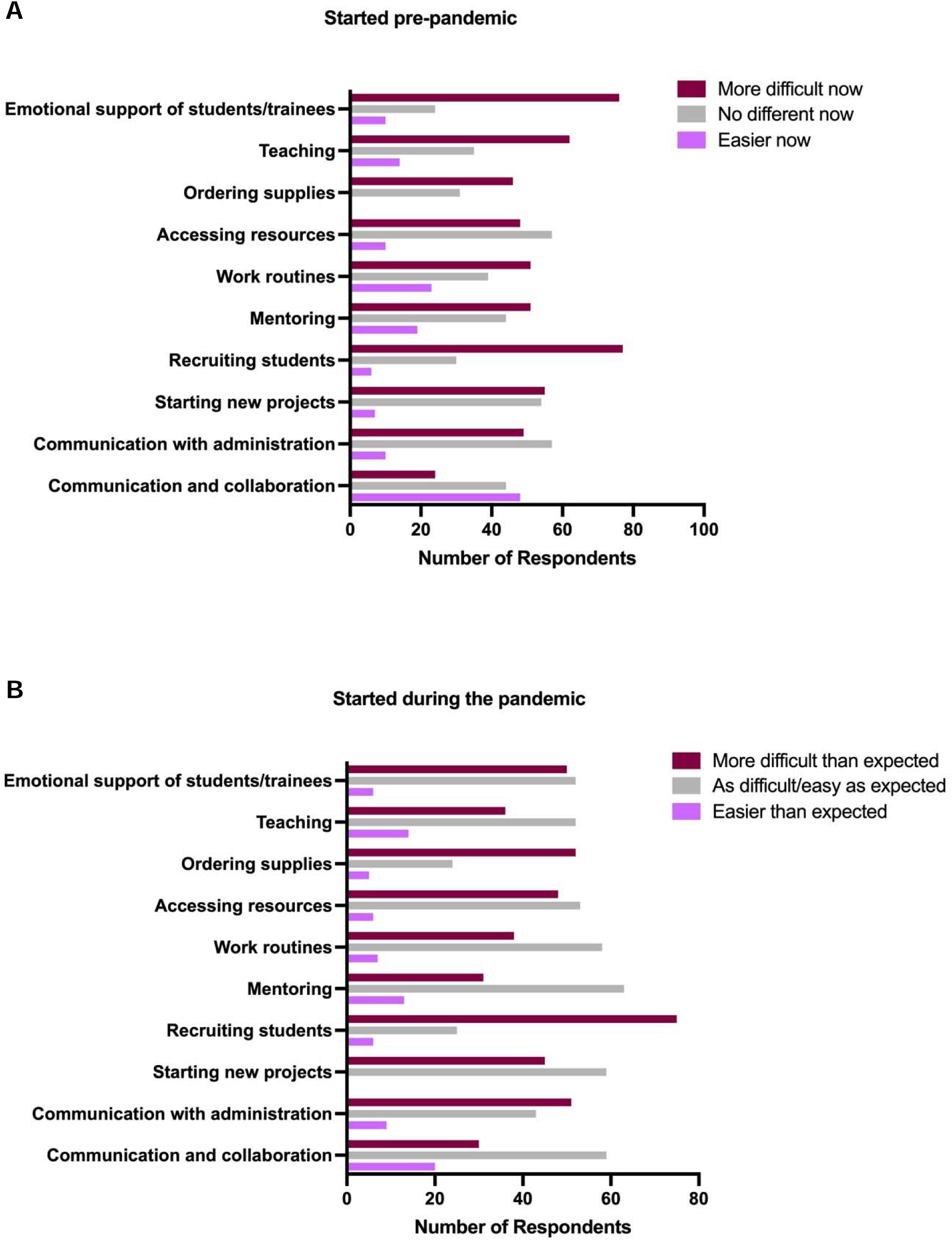
Assessment of work duties by (**A**) faculty who started before the pandemic, and (**B**) faculty who started during the pandemic.

### Burnout inventory

We found that 62% (95% confidence interval [95%CI]: 56-67%) of respondents met the criteria for overall high burnout, 64% (95%CI: 57-70%) met the criteria for high exhaustion, and almost all respondents met criteria for high disengagement (91%, 95%CI: 87-95%, Figure 4). Given the high prevalence of these outcomes, it was not possible to examine the relationship between burnout and other demographic variables (e.g., gender, career stage, etc.). Nevertheless, we note that increased number of years as an independent researcher was significantly associated with decreased odds of high disengagement (odds ratio=0.67, 95%CI 0.50-0.91, Table 6).

**Figure 4:**
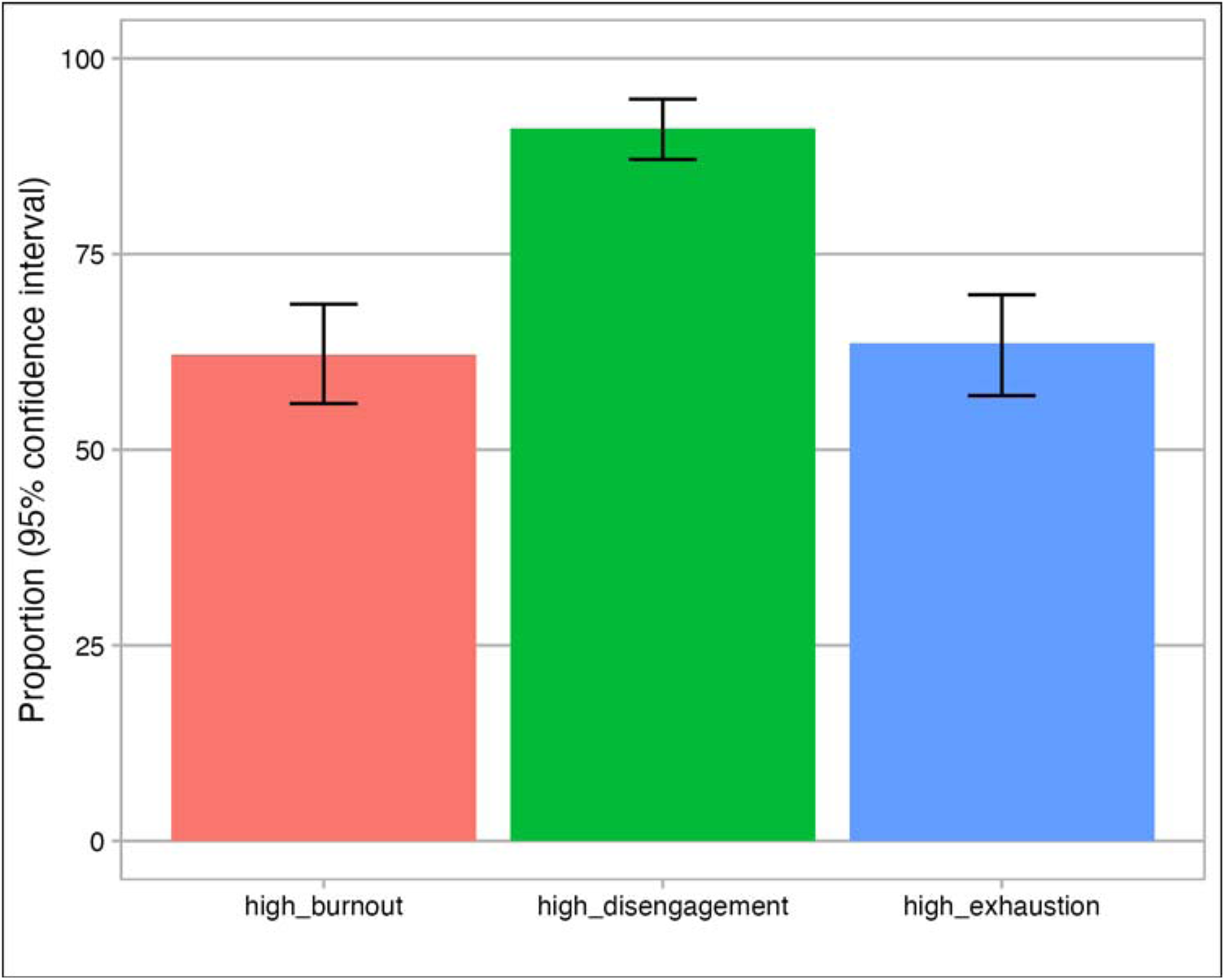
Proportion of early career health researchers in Canada meeting criteria for high burnout, disengagement or exhaustion using validated measure of burnout.

**Table 6:** Univariate regression analyses for the association between various characteristics with burnout domains and subdomains (n=225).

|  | High Burnout | High Disengagement | High Exhaustion |
| --- | --- | --- | --- |
|  | Odds ratio (95%CI) | Odds ratio (95%CI) | Odds ratio (95%CI) |
| **Age group** |  |  |  |
| 31-35 | Reference | Reference | Reference |
| 36-40 | 0.87 (0.37-2.04) | 1.37 (0.25-7.45) | 0.94 (0.40-2.23) |
| 41-45 | 0.75 (0.30-1.86) | 0.44 (0.09-2.20) | 0.80 (0.32-1.99) |
| >45 | 0.68 (0.23-2.02) | 0.86 (0.11-6.56) | 0.68 (0.23-2.02) |
| **Gender identity** |  |  |  |
| Man | Reference | Reference | Reference |
| Woman, non-binary or other | 0.58 (0.32-1.05) | 0.93 (0.36-2.58) | 0.56 (0.31-1.02) |
| **Years as independent researcher** |  |  |  |
| Per year (from 1-7) | 0.94 (0.80-1.11) | **0.67 (0.50-0.91)** | 0.95 (0.81-1.12) |
| **Caregiver of children** |  |  |  |
| No | Reference | Reference | Reference |
| Yes | 0.94 (0.54-1.65) | 1.03 (0.39-2.74) | 0.95 (0.54-1.67) |

95%CI: 95% confidence interval

### Qualitative

Our survey also provided Canadian ECHRs the opportunity to share their experience as an early career investigator during the pandemic. Three themes within this qualitative dataset emerged: ongoing benefits/problems preceding the COVID-19 pandemic; unintended outcomes of strategies to manage/prevent/contain COVID-19; and reasons to stay in their current position (Figure 5, Table 7a).

**Figure 5:**
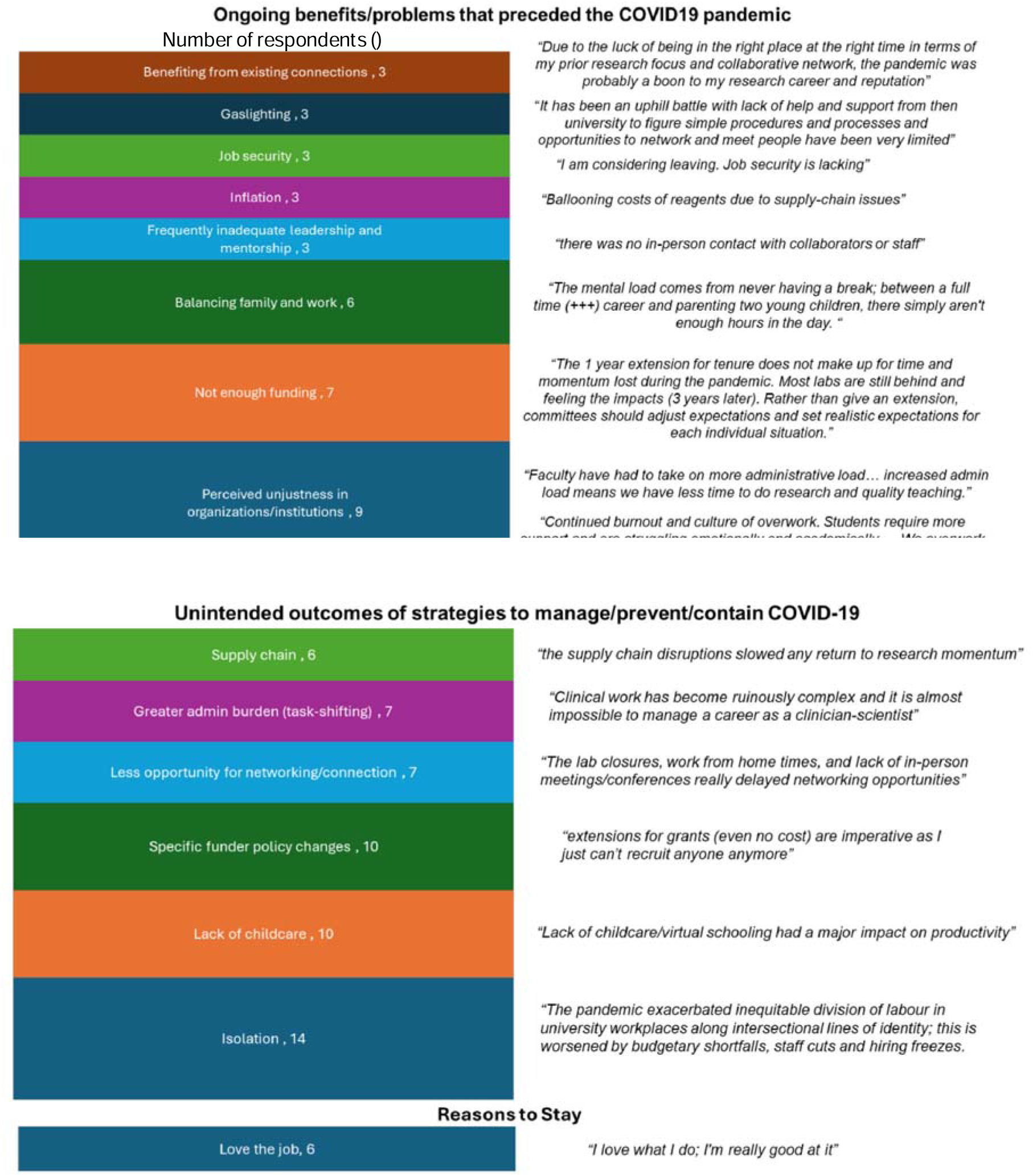
Thematic analysis to identify themes related to early career researchers’ experiences during the COVID-19 pandemic, along with sample quotes from individuals. These results are also presented in Table 7.

**Table 7:** Thematic analysis of respondents’ experiences as an early career researcher during the COVID-19 pandemic.

| **A) Identified themes** |  |  |
| --- | --- | --- |
| **Ongoing problems that preceded the COVID19 pandemic** | **Unintended outcomes of strategies to manage/prevent/contain COVID-19** | **Reasons to stay** |
| Perceived unjustness in organizations/institutions (9)<br>Not enough funding (7)<br>Balancing family and work (6)<br>Frequently inadequate leadership and mentorship (3)<br>Inflation (3)<br>Job security (3)<br>Gaslighting (3)<br>Benefiting from existing connections (3) | Isolation (14)<br>Lack of childcare (10)<br>Specific funder policy changes (10)<br>Less opportunity for networking/connection (7)<br>Greater admin burden (task-shifting) (7)<br>Supply chain (6) | Love the job (6) |

| **B) COVID-19 related outcomes** |  |  |  |
| --- | --- | --- | --- |
| **Research-related** | **Student-related** | **Personal** | **Broader** |
| Decreased productivity (21)<br>Ongoing impacts on professional network and connections (10)<br>Compromised work (5)<br>Success for funding apps (4) | Impaired recruitment of students (5)<br>Teaching related (3) | Low moral/discouraged (17)<br>Burnout (8)<br>Intent to leave (5) | Increased convenience/hybrid work (7)<br>Pool/training of future researchers (6)<br>Inequality of outcomes (e.g., senior vs. junior researchers) (4) |

Note: The same results presented in Panel A are presented in Figure 5.

Multiple issues were seen as ongoing problems preceding the COVID-19 pandemic, most prominently: perceived injustices within institutions, inadequate funding, and difficulties balancing work and family life. In relation to fairness within the institution, one respondent indicated that the “pandemic exacerbated inequitable division of labor in university workplaces along intersectional lines of identity” and that this was “worsened by budgetary shortfalls, staff cuts and hiring freezes.” Issues with the balance of work and family life were brought into particular focus by a respondent who noted the “mental load comes from never having a break; between a full time (+++) career and parenting two young children, there simply aren’t enough hours in the day.”

Isolation was the most frequently reported unintended outcome of strategies implemented to prevent or contain COVID-19 (Figure 5, Table 7a). Work buildings were described as deserted and respondents struggled to find people to ask, “simple questions on campus” to, such as “how does the printer work? where do people print posters on campus?”. The lack of childcare and school closures also significantly impacted the lives of ECHRs with pre-school and school-aged children. As one respondent noted, “as a parent of two young children, the caregiving management, even once childcare facilities opened up again was significantly higher.” Although respondents recognized that policies had been adjusted to mitigate pandemic-related impacts on academic careers, many felt these adjustments were mis-judged or not carefully implemented. As one respondent stated: “Although funding agencies have made room to explain delays related to COVID-19, for national funding opportunities, the Canadian Common CV does not allow us to report our contributions beyond the past 5 years…We can expect this 5-years restriction to disproportionately affect ECRs who were most impacted negatively by COVID19.”

The final theme of reasons to stay provides necessary context, especially when considering the preponderance of negative work-related experiences, feelings and impacts shared by respondents (Figure 5, Table 7a). Multiple respondents took the opportunity to share their love for health research and to recognize their privileged position as a researcher in a position to conduct independent research. As one respondent shared, “what keeps me motivated is my research, the communities I partner with, the participants, the innovative work, and my students.”

Beyond these themes, respondents also described outcomes related to COVID-19 and its management that included research-related, student-related, personal and broad (Table 7b). The most-reported outcomes of COVID-19 were research-related (coded 40 times within the 80 responses). Decreased productivity was the most frequently described COVID-19 research-related outcome. As one respondent noted, “because data collection was so slow for the first two years, the lab’s first papers have been delayed. This has resulted in less preliminary data for grants and several grant review comments on my lab’s lack of productivity.”

In-line with our quantitative findings, impaired student recruitment was highlighted frequently. One respondent noted “student recruitment and retention during the pandemic was difficult. International students could not enter the country, or had key funding cut by provincial agencies which caused them to consider withdrawing from the program. Local students had delays in graduation which presented as a gap in students looking for graduate opportunities.”

Personal outcomes related to the COVID-19 pandemic also appeared prominently in our findings, with many reporting low morale and burnout. As one respondent noted, “the success rate for grants in the country are so low that it’s almost not worth applying, especially in some disciplines (like nursing) where the teaching loads and family factors are incredibly high. A lot of my colleagues give up.” The term burnout was explicitly used in multiple responses and this quote captured the essence of many other responses in the category of personal outcomes of the COVID-19 pandemic: “I love what I do; I’m really good at it. And, I don’t know if this is sustainable.”

Broader outcomes of COVID-19 were those that were perceived to extend beyond the individual institution of employment and the respondent’s personal life. For example, the increased convenience of remote and hybrid work was seen as an outcome with both positive and negative effects. An advantage mentioned by a respondent was “… that some things became a bit more convenient, specifically the increase in normalizing meeting remotely (via Zoom) has made a lot of meetings more flexible and much easier to schedule.” Less positively, another respondent noted that it had “been difficult to juggle responsibilities in an on-line environment.”

## Discussion

Health knowledge is essential to supporting evidence-based healthcare practice^13,14^ and a pool of engaged and productive ECHRs is necessary for a strong and sustainable health research ecosystem. Given that the costs of faculty turnover are high, ^14^ we aimed to map the demographics and understand the experience of Canadian ECHRs in the context of the COVID-19 pandemic. We found that survey respondents were a diverse group of researchers that was representative of ECHRs applying to the largest Canadian health funding agency. Importantly, the characteristics of Canadian ECHRs mirrors the demographics of early career researchers in other countries including the USA,^15^ Australia^16^, Sweden^17^, and the United Kingdom ^18^ suggesting that our findings may reflect the situation beyond the Canadian health research ecosystem.

Most troublingly, our survey revealed that Canadian ECHRs exhibited high levels of burnout and appeared to be struggling following the COVID-19 pandemic. Over 60% of the ECHR survey respondents met the criteria for high burnout, indicating that the sustainability of the ECHR workforce in Canada is in jeopardy. Burnout is a growing issue in the modern workforce and is known to negatively impact work satisfaction and employee retention.^19,20^ Almost all respondents met the criteria for high disengagement, a problematic finding given that ECHRs represent the future of health research and innovation. Studies in the USA and Australia report similar levels (55-64%) of burnout (using self-report burnout) among faculty,^21,22^ suggesting that this phenomenon may be widespread. Similarly, a meta-analysis of burnout among public health professional, including academics, revealed a pooled proportion of burnout of 39%, although this ranged from 10-85% across individual studies.^19^ Importantly, when studies conducted before and during the COVID-19 pandemic were analyzed separately, an increase of burnout from 39% to 42% was observed.^3,7^ Although there has been significant media coverage of burnout in health academics, particularly within the context of the COVID-19 pandemic,^23,24^ its short and long-term impacts on ECHRs remain unclear.

Addressing burnout among ECHRs is critical. Burnout, defined as emotional exhaustion and cynicism, among Canadian nursing faculty at all career stages was associated with lower career satisfaction and an intention to leave their employment.^25,26^ Troublingly, attrition related to the unsustainability of academic careers further exacerbates factors contributing to burnout for those that remain in the job. Consistently, working in under-resourced nursing faculties and departments was found to increase the likelihood that faculty will seek out alternative employment, whether in other academic institutions or outside of academia. For many PhD-trained health researchers, jobs in industry are particularly attractive given that they generally offer higher compensation.^26,27^

Our survey highlighted several challenges faced by ECHRs because of the COVID-19 pandemic. The inability to form professional connections and networks emerged as a key challenge in both our quantitative and qualitative results. Several studies have emphasized the importance of supportive work environments in the success of faculty.^14,26^ Although support with research activities (e.g. with grant writing) from administrators and those in leadership positions has been reported as particularly important to the productivity, success, and retention of ECHRs, our respondents noted that obtaining such support was particularly difficult.

Several institutions and funding agencies implemented strategies to support ECHRs during the COVID-19 pandemic, signaling that they acknowledged that this group of researchers were disproportionately affected by this global event. Nevertheless, it is not clear if these strategies were effective. Among our respondents, the uptake or intent to request COVID-19-related extensions to the tenure clock was only one-quarter of those offered the extensions. Poor uptake may be linked to the fact that tenure is often associated with higher compensation (when coupled with promotion), and in many cases, necessary for continued employment. Consistently, other extensions to the tenure clock, such as those due to parental leaves, have been reported to ineffectively equalize assessments of productivity for researchers.^28^ Instead, “net academic age,” is emerging as a more nuanced means to account for factors that can impact academic productivity.

The COVID-19 pandemic may have amplified existing challenges for ECHRs, and particularly affected those with increased domestic and caregiving responsibilities, which are disproportionally women.^2^ As a result, many ECHRs experienced isolation, frustration, and decreased productivity.^4,5^ In some instances, the pandemic’s mental toll was further exacerbated by inadequate institutional support. ECHRs encountered delays in their projects, gaps in funding, and uncertainty regarding their career prospects. While offering some benefits, the shift to virtual conferences failed to provide the same collaborative opportunities as in-person networking.^6^ As the pandemic continued, many ECHRs began to reflect on the importance of work-life balance, increasing the emphasis on family and personal well-being.^5^

## What can be done

The levels of burnout in our sample of Canadian ECHRs is concerning; burnout is significantly impacted by organizational climate and culture emphasizing the need for action at various levels.^25^ A structural rethink of how academic institutions are organized, how they measure performance, and how they recognize the value of faculty may be required to achieve a consequential reduction in rates of burnout and ultimately attrition among ECHRs.^29^ Findings from a 2021 study of Canadian nursing faculty support the need for comprehensive resources and initiatives to better support early career researchers with their research programs.^25^ Retention of faculty was improved following targeted initiatives aimed at enhancing faculty’s capacity to achieve work-life balance, such as by setting a maximum for hours worked per week and instituting flexible practices in the workplace.^13^ Leaders across many academic settings should establish “manageable” expectations, promote and model work-life balance^22,25^ and value the quality of work over quantity.^25^

## Strengths and Limitations

Our survey had strengths which included a large population of Canadian ECHR respondents that is representative of the ECHRs that apply to the national health research funding agency. Nevertheless, reliance on a convenience sample meant that we only captured a proportion of all Canadian ECHRs – exactly what proportion of all ECHRs in Canada we captured cannot be calculated with available data. French-only speaking ECHRs were likely excluded since our survey was only available in English. Our survey also failed to collect information on Indigenous identity data because we were unable to identify Indigenous scholars with the capacity to engage with the project.

## Conclusions

We have reported on the first survey of a broad sample of Canadian ECHRs, providing baseline information that can be used to inform, develop, implement, and evaluate policies aimed at supporting ECHRs in Canada. Our survey results suggest that Canadian ECHRs share characteristics with early career researchers in other countries, suggesting that successful policies implemented in other countries to support ECHRs may also work in Canada. Supporting ECHRs is critical to the flourishing of the Canadian health research ecosystem. Understanding Canadian ECHRs and their job experiences is an important first step in developing strategies to engage and retain this important group of health researchers.

## Declarations

### Ethics approval and consent to participate

The protocol was reviewed to be in accordance with ethical standards for research involving humans (University of Prince Edward Island certificate #16778). Participants were presented with a letter of information prior to beginning the study and were informed that their responses would be anonymous and that they would be unable to withdraw their responses once they had completed the survey.

### Availability of data and materials

De-identified data could be made available to interested researchers by contacting the authors and with necessary ethical approval and data sharing agreements. Researchers interested in addressing research questions related to grant funding can contact the CIHR at.

### Competing interests

All authors declare no conflicts related to this work. The views expressed in this Article are those of the authors and do not necessarily reflect those of the CIHR or the Government of Canada.

### Funding

This work was funded in part by a CIHR Planning and Dissemination grant (NPAs: Hewko, Kowalec, award number 486136). GEL holds the Canada Research Chair in Adipocyte Development.

## Authors’ contributions

SH, KK, EEM, and LA analyzed and interpreted data. All authors apart from LD and GL contributed to acquiring funding for the project. All authors were involved in preparing the initial manuscript draft and read and approved the submitted manuscript.

## Supporting information

Copy of survey

## Acknowledgements

We thank Professor Liisa Galea and Professor Jim Woodgett for their valuable input in designing the survey. We thank Freda Warner and Audrey Arnot (CIHR Funding Analytics staff) for extracting the dataset. The CIHR is committed to collaborating with the research community to advance knowledge on best practices in peer review without unduly influencing the conclusions drawn by its collaborators. For this reason, CIHR employees are not currently permitted to be co-authors on articles describing analyses of the agency’s programs. Without the active participation of these CIHR employees and their contributions, this article would not have been possible.

## Notes

### Competing Interest Statement

The authors have declared no competing interest.

