## Supplementary Tables for "Characteristics of early career health researchers and experiences of burnout during the COVID-19 pandemic in Canada"

12: Cardiometabolic axis, Centre de Recherche du Centre hospitalier de l'Université de Montréal (CRCHUM), Canada

13: Department of Cellular and Molecular Medicine, University of Ottawa, Canada

14: Faculty of Kinesiology & Recreation Management, University of Manitoba, Canada

15: The Children’s Hospital Research Institute of Manitoba (CHRIM), Canada

16: School of Library and Information Science. Université de Montréal, Canada

17: Centre de recherche en santé publique (CReSP), Canada

18: Division of Pulmonary Medicine, Faculty of Medicine and Dentistry, University of Alberta, Canada

*Shared first-author

**Table S1:** Job characteristics and demographics of Canadian early career health researchers, by institute type

|  | **Institution type*** | | | |
| --- | --- | --- | --- | --- |
|  | **Comprehensive** | **Medical/ doctoral** | **Hospital** | **Primarily undergraduate** |
| **N** | 78 (34.6) | 79 (35.1) | 41 (18.2) | 21 (9.3) |
| **Age, years** | 40 (37, 42) | 39 (37, 42.5) | 38 (35, 42) | 40 (38, 43) |
| **Gender Identity** |  |  |  |  |
| Woman | 53 (67.9) | 42 (53.2) | 17 (41.4) | 6 (28.5) |
| Man | 21 (26.9) | 32 (40.5) | 24 (58.5) | 16 (76.2) |
| Mix of gender identities | ¶ | ¶ | 0 | 0 |
| Prefer not to answer | ¶ | ¶ | 0 | 0 |
| **Years in independent position** | 3 (2, 5) | 4 (3, 5) | 3 (3, 4) | 4 (3, 5) |
| **Province or territory**** |  |  |  |  |
| Alberta | ¶ | 11 (13.9) | 0 | ¶ |
| British Columbia | 15 (19.2) | 7 (8.8) | ¶ | ¶ |
| Prairies | 7 (8.9) | 7 (8.8) | ¶ | ¶ |
| Atlantic Canada | ¶ | 6 (7.6) | 0 | 5 (22.7) |
| Ontario | 28 (35.9) | 23 (29.1) | 20 (48.7) | 7 (31.8) |
| Québec | 16 (20.5) | 21 (26.5) | 18 (43.9) | ¶ |
| **Current rank** |  |  |  |  |
| Assistant Professor | 63 (80.8) | 60 (75.9) | 34 (82.9) | 19 (86.4) |
| Associate Professor | 13 (16.7) | 18 (22.8) | 5 (12.2) | ¶ |
| Adjunct | ¶ | ¶ | ¶ | 0 |
| **Research pillar** |  |  |  |  |
| Biomedical | 46 (58.9) | 38 (48.1) | 9 (21.9) | 14 (66.7) |
| Clinical | 10 (12.8) | 14 (17.7) | 16 (39.0) | ¶ |
| Health services | 9 (11.5) | 15 (18.9) | 7 (17.1) | ¶ |
| Social, cultural, environmental, and population health | 13 (16.7) | 11 (13.9) | 9 (21.9) | ¶ |
| **Job characteristics** |  |  |  |  |
| Teaching stream | 5 (6.4) (n = 78) | 8 (10.1) | 7 (17.1) | ¶ |
| Clinical stream | ¶ | 27 (34.2) | 17 (41.5) | ¶ |
| **Grant agencies applied to** |  |  |  |  |
| Tri-agency | 72 (92.3) | 76 (96.2) | 35 (85.4) | 18 (81.2) |
| Federal agency | 36 (46.2) | 50 (63.2) | 14 (34.1) | 9 (40.9) |
| Non-profit/charity | 46 (58.9) | 59 (74.6) | 28 (68.3) | 10 (45.5) |
| Province/territory | 50 (64.1) | 62 (78.5) | 27 (65.9) | 7 (31.8) |
| US agency | 23 (29.4) | 21 (26.5) | 13 (31.7) | 5 (22.8) |
| International not US | 8 (10.3) | 18 (22.7) | 7 (17.0) | 0 |

Numbers are reported as n (%) or median (interquartile range). ¶ = Suppressed as N<5. *N<5 persons reported their institution type as "other" or "government agency". **We had no respondents from any Canadian territories.

**Table S2:** Job characteristics and demographics of Canadian early career health researchers, by gender identity

|  | **Gender identity*** | |
| --- | --- | --- |
|  | **Woman** | **Man** |
| **N** | 139 (62) | 77 (34) |
| **Age, years** | 40 (37, 43) | 38 (36, 42) |
| **Independent position, years** | 3 (2, 5) | 4 (3, 5) |
| **Province or territory**** |  |  |
| Alberta | 8 (5.7) | 8 (10.4) |
| British Columbia | 20 (14.4) | 4 (5.2) |
| Prairies | 18 (12.9) | 7 (9.1) |
| Atlantic Canada | 9 (6.5) | 5 (6.5) |
| Ontario | 45 (32.4) | 33 (42.8) |
| Québec | 39 (28.1) | 20 (25.9) |
| **Current rank** |  |  |
| Assistant Professor | 116 (83.5) | 58 (75.3) |
| Associate Professor | 20 (14.4) | 16 (20.8) |
| Adjunct | 3 (2.16) | 3 (3.90) |
| **Research pillar** |  |  |
| Biomedical | 54 (38.8) | 50 (64.9) |
| Clinical | 28 (20.1) | 12 (15.6) |
| Health services | 25 (18.1) | 9 (11.7) |
| Social, cultural, environmental, and population health | 32 (23.0) | 5 (6.5) |
| **Institute type** |  |  |
| Comprehensive | 53 (38.1) | 21 (27.3) |
| Medical/doctoral | 42 (30.2) | 32 (41.6) |
| Hospital | 24 (17.3) | 17 (22.1) |
| Primarily undergraduate | 16 (11.5) | 6 (7.8) |
| Government | ¶ | ¶ |
| Other | ¶ | ¶ |
| **Job characteristics** |  |  |
| Teaching stream | 12 (8.6) | 10 (13.0) |
| Clinical stream | 21 (15.1) | 16 (20.8) |
| **Grant agencies applied to** |  |  |
| Tri-agency | 126 (90.6) | 69 (89.6) |
| Federal agency | 64 (46.0) | 44 (57.1) |
| Non-profit/charity | 82 (58.9) | 59 (76.6) |
| Province/territory | 89 (64.0) | 55 (71.4) |
| US agency | 34 (24.5) | 23 (29.9) |
| International not US | 16 (11.5) | 13 (16.9) |

Numbers are reported as n (%) or median (interquartile range). ¶ = Suppressed as N<5. *N<5 persons reported their institution type as "other" or "government agency". **We had no respondents from any Canadian territories.

**Table S3:** Reasons for not applying for grants by early career health researchers, by overall sample and by research pillar

|  | **Total** | **Research Pillar** | | | |
| --- | --- | --- | --- | --- | --- |
|  |  | **Biomedical** | **Clinical** | **Health services** | **Social, cultural, environmental, and population health** |
| **N** | 225 | 109 (48.4) | 41 (18.2) | 36 (16) | 38 (16.9) |
| **Reasons for not applying for as many grants as principal investigator as expected** | | | | | |
| On leave | 26 (11.5) | 7 (6.4) | 6 (14.6) | 6 (16.7) | 7 (18.4) |
| Already sufficiently funded | 42 (18.6) | 18 (16.5) | 5 (12.2) | ¶ | 16 (42.1) |
| Not ready | 66 (29.3) | 39 (35.8) | 11 (26.8) | 11 (30.6) | 5 (13.2) |
| Discouraged | 65 (28.8) | 26 (23.9) | 13 (31.7) | 14 (38.9) | 11 (28.9) |
| Inadequate resources | 89 (39.5) | 35 (32.1) | 17 (41.5) | 19 (52.8) | 17 (44.7) |
| Setting up lab | 36 (16.0) | 28 (25.7) | ¶ | ¶ | ¶ |
| Applied for as many as expected | 64 (28.4) | 36 (33) | 12 (29.3) | 5 (13.9) | 11 (28.9) |
| **Reasons for not applying for as many grants as co-investigator as expected** | | | | | |
| On leave | 13 (5.7) | 5 (4.5) | ¶ | ¶ | ¶ |
| Already sufficiently funded | 15 (6.7) | 9 (8.3) | ¶ | ¶ | ¶ |
| Not ready | 29 (12.8) | 20 (18.3) | ¶ | ¶ | ¶ |
| Discouraged | 32 (14.2) | 6 (5.5) | 7 (17.1) | 11 (30.6) | 8 (21.1) |
| Inadequate resources | 44 (19.6) | 19 (17.4) | 7 (17.1) | 11 (30.6) | 7 (18.4) |
| Insufficient data | 29 (12.9) | 21 (19.3) | ¶ | ¶ | ¶ |
| Applied for as many as expected | 107 (47.5) | 47 (43.1) | 22 (53.7) | 16 (44.4) | 21 (55.3) |

Numbers are reported as n (%) or median (interquartile range), ¶ = Suppressed as N<5.

**Table S4:** COVID-19 related descriptives by gender identity and institution type

|  | **Gender identity** | | **Institution** | | | |
| --- | --- | --- | --- | --- | --- | --- |
|  | **Woman** | **Man** | **Comprehensive** | **Medical/ doctoral** | **Hospital** | **Primarily undergraduate** |
| **N** | 139 (61.7) | 77 (34.2) | 78 (34.7) | 79 (35.1) | 41 (18.2) | 21 (9.8) |
| **COVID-19 time extension for tenure** | | | | | | |
| Yes | 73 (52.5) | 34 (44.2) | 50 (64.1) | 44 (55.6) | ¶ | 16 (76.2) |
| No | 20 (14.4) | 15 (19.5) | 10 (12.8) | 10 (12.7) | 14 (34.1) | ¶ |
| University does not have tenure track | 7 (5.0) | ¶ | ¶ | ¶ | 8 (19.5) | 0 |
| I don’t know | 38 (27.3) | 21 (27.3) | 0 | 21 (26.5) | 14 (34.1) | ¶ |
| Prefer not to answer | ¶ | ¶ | 17 (21.8) | ¶ | ¶ | 0 |
| **COVID-19 length of extension (n = 114)** | | | | | | |
| 1 year | 64 (87.6) | 31 (91.2) | 45 (90.0) | 39 (88.9) | ¶ | 15 (93.8) |
| 2 years | 9 (12.3) | ¶ | 5 (10.0) | 5 (11.1) | ¶ | ¶ |
| **COVID-19 tenure extension used (n = 113)** | | | | | | |
| Definitely no (or didn’t) | 26 (35.6) | 20 (58.8) | 20 (40.0) | 19 (43.4) | 0 | 10 (62.5) |
| Probably no | 12 (16.4) | 6 (17.6) | 10 (20.0) | 8 (18.6) | 0 | ¶ |
| Haven’t decided yet | 14 (19.1) | 0 | 9 (18.0) | ¶ | 0 | ¶ |
| Probably yes | 5 (6.8) | ¶ | 6 (12.0) | ¶ | ¶ | 0 |
| Definitely yes | 15 (20.5) | 6 (17.6) | 5 (10.0) | 12 (27.2) | ¶ | ¶ |
| **Impact of citizenship status on ECHR status during COVID-19 (n = 219)** | | | | |  |  |
| Impacted my experience very negatively | 6 (4.32) | 8 (10.4) | 7 (8.9) | ¶ | ¶ | ¶ |
| Impacted my experience negatively | 19 (13.7) | ¶ | 13 (16.7) | 8 (10.1) | ¶ | ¶ |
| Had no impact on my experience | 99 (71.2) | 64 (83.1) | 51 (65.4) | 62 (78.5) | 33 (80.5) | 18 (81.2) |
| Impacted my experience positively | 7 (5.0) | 0 | ¶ | 0 | ¶ | 0 |
| Impacted my experience very positively | ¶ | 0 | ¶ | 0 | ¶ | ¶ |
| **Mental health changes during COVID-19** | | | |  |  |  |
| Significantly worsened | 18 (12.9) | 15 (19.5) | 12 (15.4) | 17 (21.5) | 6 (14.6) | 0 |
| Moderately worsened | 75 (54.0) | 30 (39) | 42 (53.9) | 35 (44.3) | 16 (39.0) | 14 (63.6) |
| Has not changed | 40 (28.8) | 26 (33.8) | 21 (26.9) | 20 (25.3) | 18 (43.9) | 7 (31.8) |
| Moderately improved | ¶ | 6 (7.8) | ¶ | 5 (6.3) | ¶ | ¶ |
| Improved significantly | ¶ | 0 | 0 | 0 | 0 | 0 |
| Prefer not to answer | ¶ | 0 | 0 | ¶ | 0 | 0 |
| **Pivoted to COVID-19 research** | | |  |  |  |  |
| No | 60 (43.2) | 48 (62.3) | 39 (50.6) | 43 (54.4) | 15 (36.5) | 12 (57.1) |
| Partially | 73 (52.5) | 24 (31.2) | 37 (48.1) | 31 (39.2) | 24 (58.5 | 8 (38.1) |
| Fully | 6 (4.3) | ¶ | ¶ | ¶ | ¶ | ¶ |

Numbers are reported as n (%) or median (interquartile range); ¶ = Suppressed as N<5.

**Table S5:** Reasons for not applying for grants by early career health researchers, by gender identity and institution type

|  | **Gender identity** | | **Institution** |  |  |  |
| --- | --- | --- | --- | --- | --- | --- |
|  | **Woman** | **Man** | **Comprehensive** | **Medical/ doctoral** | **Hospital** | **Primarily undergraduate** |
| **Reasons for not applying for as many principal investigator grants as expected** | | | | | |  |
| On leave | 21 (15.1) | ¶ | 7 (8.9) | 7 (8.8) | 7 (17.1) | 5 (23.8) |
| Already sufficiently funded | 31 (22.3) | 10 (13.0) | 15 (19.2) | 13 (16.5) | 5 (12.2) | 8 (38.1) |
| Not ready | 45 (32.4) | 21 (27.3) | 23 (29.5) | 20 (25.3) | 12 (29.3) | 9 (42.8) |
| Discouraged | 38 (27.3) | 26 (33.8) | 19 (24.3) | 24 (30.3) | 14 (34.2) | 6 (28.5) |
| Inadequate resources | 56 (40.3) | 31 (40.3) | 36 (46.2) | 27 (34.2) | 18 (43.9) | 5 (23.8) |
| Setting up lab | 22 (15.8) | 14 (18.2) | 17 (21.7) | 12 (15.3) | 5 (12.2) | ¶ |
| Applied for as many as expected | 39 (28.1) | 22 (28.6) | 24 (30.7) | 23 (29.1) | 11 (26.8) | 5 (23.8) |
| **Reasons for not applying for as many co-investigator grants as expected** | | | | | |  |
| On leave | 11 (7.9) | ¶ | 5 (6.4) | ¶ | ¶ | 0 (0) |
| Already sufficiently funded | 10 (7.2) | ¶ | 6 (7.6) | ¶ | ¶ | ¶ |
| Not ready | 20 (14.4) | 9 (11.7) | 10 (12.8) | 9 (11.4) | ¶ | 5 (23.8) |
| Discouraged | 20 (14.4) | 12 (15.6) | 8 (10.3) | 11 (13.9) | 8 (19.5) | ¶ |
| Inadequate resources | 28 (20.1) | 16 (20.8) | 21 (26.9) | 10 (12.7) | 9 (21.9) | ¶ |
| Insufficient data | 22 (15.8) | 7 (9.1) | 13 (16.7) | 9 (11.4) | ¶ | ¶ |
| Applied for as many as expected | 68 (48.9) | 34 (44.2) | 38 (48.7) | 37 (46.8) | 20 (48.8) | 10 (47.6) |

Numbers are reported as n (%) or median (interquartile range); ¶ = Suppressed as N<5.
