## Supplementary material for "Characteristics of early career health researchers and experiences of burnout during the COVID-19 pandemic in Canada": Copy of survey

**Principal Investigators**:

Sarah Hewko, RD PhD, Assistant Professor, Department of Applied Human Sciences, University of Prince Edward Island

Kaarina Kowalec, MSc. PhD, Co-Chair of the Association for Canadian Early Career Health Researchers and Assistant Professor of Pharmacy, University of Manitoba

**STUDY INFORMATION**

**Invitation to Participate**

You are invited to participate in a research study designed to collect information on Canadian Early Career Health Researchers (ECHRs): (1) characteristics associated with diversity, (2) funding history, (3) COVID-19 related impacts on career and (4) general feelings about work. We are hoping for a minimum of 200 respondents.

**Purpose of the Study**

We anticipate that our study findings will gain national attention and be of direct, practical use to university administrators, health research institutions, and decision-makers at health research funding agencies across Canada. Supporting ECHRs to conduct research in health and protecting their ability to thrive in the Canadian research ecosystem: i) ensures that Canadians have access to context-specific, evidence-informed health care, ii) prevents attrition of highly-trained personnel via emigration, and ii) enhances the quality of education provided within Canada’s post-secondary institutions.

**Inclusion Criteria**

You are eligible to participate if you are currently an independent health researcher at a Canadian university or research institute and are still considered an “early career” scientist (with consideration for eligible delays such as parental leaves and COVID-19 extensions). For eligibility purposes, we are employing the Canadian Institutes for Health Research definitions for both researcher (independent) and researcher (early career) (https://cihr-irsc.gc.ca/e/34190.html#r).

**Study Procedures**

By continuing from this page onto the survey, you are consenting to participate in this study. We expect the survey to take you no more than 20 minutes to complete. You may want to pull up your curriculum vitae to reference for questions about grant applications. After we have confirmed your eligibility to participate, you are free skip questions you would prefer not to answer. You can withdraw from the study at any time by closing this tab in your browser. You may stop completing the survey at any time without repercussions. Study data will be stored indefinitely on a password-protected, encrypted University of Prince Edward Island server.

**Possible Risks and Harms**

There are no known or anticipated risks associated with participating in this study. You are not being asked to waive any of your rights. You can decline to answer any question(s). If you experience any distressing emotions following completion of this study, you can contact Wellness Together Canada at 1-866-585-0445 or text WELLNESS to 741741.

**Possible Benefits**

We cannot guarantee any direct benefits because of your participation in this study. However, it is our hope that our study findings will gain national attention and be of direct, practical use to university adminstrators, health research institutions, and decision-makers at health research funding agencies across Canada. Supporting ECHRs to conduct research in health and protecting their ability to thrive in the Canadian research ecosystem: i) ensures that Canadians have access to context-specific, evidence-informed health care, ii) prevents attrition of highly-trained personnel via emigration, and ii) enhances the quality of education provided within Canada's post-secondary institutions.

**Confidentiality and Privacy**

Your survey responses are entered anonymously. For this reason, you cannot withdraw your data from analysis once you have submitted your survey responses. All reporting will be at the aggregate level, such that no one can be identified based on disseminated results/findings.

**Contacts for Study Questions or Problems**

If you require any further information regarding this research project or your participation in the study you may contact Sarah Hewko at (mailto:) or Kaarina Kowalec at (mailto:). If you would like to keep a copy of this information for your records, you can print this screen or take a screen shot now. This project has been reviewed by the UPEI Research Ethics Board and it complies with Tri-Council guidelines for research involving human participants. If you have any concerns about the ethical conduct of this study you can contact the UPEI Research Ethics Board at (p - (902)620-5104).

There are 50 questions in this survey.

### Demographic Information

#### Are you currently an independent health researcher (in a position where you can apply for research grants as Principal Investigator) at a Canadian university or research institute? *

❗ Choose one of the following answers

Please choose **only one** of the following:

- ◯ Yes
- ◯ No

#### Thank you for your interest in this research but you do not meet the eligibility criteria.

Only answer this question if the following conditions are met:
Answer was 'No' at question '1 [D01]' (Are you currently an independent health researcher (in a position where you can apply for research grants as Principal Investigator) at a Canadian university or research institute?)

Please write your answer here:

#### Excluding any eligible delays (e.g., parental or medical leave), how many years has it been since you were first appointed as an independent researcher? *

Only answer this question if the following conditions are met:
Answer was 'Yes' at question '1 [D01]' (Are you currently an independent health researcher (in a position where you can apply for research grants as Principal Investigator) at a Canadian university or research institute?)

❗ Choose one of the following answers
Please choose **only one** of the following:

- ◯ <1 year
- ◯ 1 year
- ◯ 2 years
- ◯ 3 years
- ◯ 4 years
- ◯ 5 years
- ◯ 6 years
- ◯ 7 years
- ◯ >7 years

#### Thank you for your interest in our research but you do not meet the criteria for eligiblity.

Only answer this question if the following conditions are met:
Answer was '>7 years' at question '3 [D02]' (Excluding any eligible delays (e.g., parental or medical leave), how many years has it been since you were first appointed as an independent researcher?)

Please write your answer here:

### In which province or territory are you employed?

Only answer this question if the following conditions are met:
Answer was 'Yes' at question '1 [D01]' (Are you currently an independent health researcher (in a position where you can apply for research grants as Principal Investigator) at a Canadian university or research institute?) *and* Answer was NOT '>7 years' at question '3 [D02]' (Excluding any eligible delays (e.g., parental or medical leave), how many years has it been since you were first appointed as an independent researcher?)

❗ Choose one of the following answers
Please choose **only one** of the following:

- ◯ Alberta
- ◯ British Columbia
- ◯ Manitoba
- ◯ New Brunswick
- ◯ Newfoundland
- ◯ Northwest Territories
- ◯ Nova Scotia
- ◯ Nunavut
- ◯ Ontario
- ◯ Prince Edward Island
- ◯ Québec
- ◯ Saskatchewan
- ◯ Yukon

### What is your current academic rank?

Only answer this question if the following conditions are met:
Answer was 'Yes' at question '1 [D01]' (Are you currently an independent health researcher (in a position where you can apply for research grants as Principal Investigator) at a Canadian university or research institute?) *and* Answer was NOT '>7 years' at question '3 [D02]' (Excluding any eligible delays (e.g., parental or medical leave), how many years has it been since you were first appointed as an independent researcher?)

❗ Choose one of the following answers
Please choose **only one** of the following:

- ◯ Assistant Professor
- ◯ Associate Professor
- ◯ Professor
- ◯ Adjunct
- ◯ Other ______

#### Which Canadian Institutes of Health Research pillar does your work primarily fall under?

Only answer this question if the following conditions are met:
Answer was 'Yes' at question '1 [D01]' (Are you currently an independent health researcher (in a position where you can apply for research grants as Principal Investigator) at a Canadian university or research institute?) *and* Answer was NOT '>7 years' at question '3 [D02]' (Excluding any eligible delays (e.g., parental or medical leave), how many years has it been since you were first appointed as an independent researcher?)

❗ Choose one of the following answers
Please choose **only one** of the following:

- ◯ Biomedical (1)
- ◯ Clinical (2)
- ◯ Health Services (3)
- ◯ Population Health (4)

### What is your current job status or situation?

Only answer this question if the following conditions are met:
Answer was 'Yes' at question '1 [D01]' (Are you currently an independent health researcher (in a position where you can apply for research grants as Principal Investigator) at a Canadian university or research institute?) *and* Answer was NOT '>7 years' at question '3 [D02]' (Excluding any eligible delays (e.g., parental or medical leave), how many years has it been since you were first appointed as an independent researcher?)

❗ Choose one of the following answers
Please choose **only one** of the following:

- ◯ Tenure track
- ◯ Tenured
- ◯ Permanent position, but neither tenured or tenure track
- ◯ Contract or term position
- ◯ Grant - tenure
- ◯ Prefer not to answer
- ◯ Other [ ]

### In which type of institution are you primarily employed?

Only answer this question if the following conditions are met:
Answer was 'Yes' at question '1 [D01]' (Are you currently an independent health researcher (in a position where you can apply for research grants as Principal Investigator) at a Canadian university or research institute?) *and* Answer was NOT '>7 years' at question '3 [D02]' (Excluding any eligible delays (e.g., parental or medical leave), how many years has it been since you were first appointed as an independent researcher?)

❶ Choose one of the following answers
Please choose **only one** of the following:

- ◯ Primarily undergraduate university
- ◯ Comprehensive university
- ◯ Medical/Doctoral university
- ◯ Government agency
- ◯ Hospital or hospital-affiliated research institute
- ◯ Prefer not to answer
- ◯ Other ____________

#### Is you position classified as teaching stream?

Only answer this question if the following conditions are met:
Answer was 'Yes' at question '1 [D01]' (Are you currently an independent health researcher (in a position where you can apply for research grants as Principal Investigator) at a Canadian university or research institute?) *and* Answer was NOT '>7 years' at question '3 [D02]' (Excluding any eligible delays (e.g., parental or medical leave), how many years has it been since you were first appointed as an independent researcher?)

❗ Choose one of the following answers
Please choose **only one** of the following:

- ◯ No
- ◯ Yes

#### Is your position classified as clinical stream?

Only answer this question if the following conditions are met:
Answer was 'Yes' at question '1 [D01]' (Are you currently an independent health researcher (in a position where you can apply for research grants as Principal Investigator) at a Canadian university or research institute?) *and* Answer was NOT '>7 years' at question '3 [D02]' (Excluding any eligible delays (e.g., parental or medical leave), how many years has it been since you were first appointed as an independent researcher?)

❗ Choose one of the following answers
Please choose **only one** of the following:

- ◯ No
- ◯ Yes

### What is the breakdown of responsibilities in your current role? (should add up to 100%)

Only answer this question if the following conditions are met:
Answer was 'Yes' at question '1 [D01]' (Are you currently an independent health researcher (in a position where you can apply for research grants as Principal Investigator) at a Canadian university or research institute?) *and* Answer was NOT '>7 years' at question '3 [D02]' (Excluding any eligible delays (e.g., parental or medical leave), how many years has it been since you were first appointed as an independent researcher?)

| | Research | Teaching | Clinical | Service | Other |
|---|---|---|---|---|---|
| **Breakdown (%)** | | | | | |

### Please indicate which term best describes your gender identity:

Only answer this question if the following conditions are met:
Answer was 'Yes' at question '1 [D01]' (Are you currently an independent health researcher (in a position where you can apply for research grants as Principal Investigator) at a Canadian university or research institute?) *and* Answer was NOT '>7 years' at question '3 [D02]' (Excluding any eligible delays (e.g., parental or medical leave), how many years has it been since you were first appointed as an independent researcher?)

❗ Choose one of the following answers
Please choose **only one** of the following:

- ◯ Gender fluid
- ◯ Man
- ◯ Nonbinary
- ◯ Trans man
- ◯ Trans woman
- ◯ Two-Spirit
- ◯ Woman
- ◯ Prefer not to answer
- ◯ Prefer to specify: ____________

#### What is your age?

Only answer this question if the following conditions are met:
Answer was 'Yes' at question '1 [D01]' (Are you currently an independent health researcher (in a position where you can apply for research grants as Principal Investigator) at a Canadian university or research institute?) *and* Answer was NOT '>7 years' at question '3 [D02]' (Excluding any eligible delays (e.g., parental or medical leave), how many years has it been since you were first appointed as an independent researcher?)

❗ Only numbers may be entered in this field.
❗ Your answer must be between 20 and 85
Please write your answer here:

#### Do you identify as a person with a physical disability?

Only answer this question if the following conditions are met:
Answer was 'Yes' at question '1 [D01]' (Are you currently an independent health researcher (in a position where you can apply for research grants as Principal Investigator) at a Canadian university or research institute?) *and* Answer was NOT '>7 years' at question '3 [D02]' (Excluding any eligible delays (e.g., parental or medical leave), how many years has it been since you were first appointed as an independent researcher?)

❗ Choose one of the following answers
Please choose **only one** of the following:

- ◯ No
- ◯ Yes
- ◯ Prefer not to answer

#### Do you identify as a person with a mental disability?

Only answer this question if the following conditions are met:
Answer was 'Yes' at question '1 [D01]' (Are you currently an independent health researcher (in a position where you can apply for research grants as Principal Investigator) at a Canadian university or research institute?) *and* Answer was NOT '>7 years' at question '3 [D02]' (Excluding any eligible delays (e.g., parental or medical leave), how many years has it been since you were first appointed as an independent researcher?)

❗ Choose one of the following answers
Please choose **only one** of the following:

- ◯ No
- ◯ Yes
- ◯ Prefer not to answer

#### Do you have any children?

Only answer this question if the following conditions are met:
Answer was 'Yes' at question '1 [D01]' (Are you currently an independent health researcher (in a position where you can apply for research grants as Principal Investigator) at a Canadian university or research institute?) *and* Answer was NOT '>7 years' at question '3 [D02]' (Excluding any eligible delays (e.g., parental or medical leave), how many years has it been since you were first appointed as an independent researcher?)

❗ Choose one of the following answers
Please choose **only one** of the following:

- ◯ No
- ◯ Yes
- ◯ Prefer not to answer

#### How many children do you have?

Only answer this question if the following conditions are met:
Answer was 'Yes' at question '1 [D01]' (Are you currently an independent health researcher (in a position where you can apply for research grants as Principal Investigator) at a Canadian university or research institute?) *and* Answer was NOT '>7 years' at question '3 [D02]' (Excluding any eligible delays (e.g., parental or medical leave), how many years has it been since you were first appointed as an independent researcher?) *and* Answer was 'Yes' at question '17 [D18]' (Do you have any children?)

❗ Only numbers may be entered in this field.
Please write your answer here:

______________________________

#### What are the ages of your children? (select all that apply)

Only answer this question if the following conditions are met:
Answer was 'Yes' at question '1 [D01]' (Are you currently an independent health researcher (in a position where you can apply for research grants as Principal Investigator) at a Canadian university or research institute?) *and* Answer was 'Yes' at question '17 [D18]' (Do you have any children?)

❗ Check all that apply
Please choose **all** that apply:

- ☐ <2 years of age
- ☐ 2-4 years of age
- ☐ 5-18 years of age
- ☐ >18 years of age

### Are you a primary caregiver for these children?

Only answer this question if the following conditions are met:
Answer was 'Yes' at question '1 [D01]' (Are you currently an independent health researcher (in a position where you can apply for research grants as Principal Investigator) at a Canadian university or research institute?) *and* Answer was NOT '>7 years' at question '3 [D02]' (Excluding any eligible delays (e.g., parental or medical leave), how many years has it been since you were first appointed as an independent researcher?) *and* Answer was 'Yes' at question '17 [D18]' (Do you have any children?)

❗ Choose one of the following answers
Please choose **only one** of the following:

◯ No
◯ Yes
◯ Prefer not to answer

### Do you have a partner?

Only answer this question if the following conditions are met:
Answer was 'Yes' at question '1 [D01]' (Are you currently an independent health researcher (in a position where you can apply for research grants as Principal Investigator) at a Canadian university or research institute?) *and* Answer was NOT '>7 years' at question '3 [D02]' (Excluding any eligible delays (e.g., parental or medical leave), how many years has it been since you were first appointed as an independent researcher?)

❗ Choose one of the following answers
Please choose **only one** of the following:

◯ No
◯ Yes
◯ Prefer not to answer

### Are you primary caregiver for:

Only answer this question if the following conditions are met:
Answer was 'Yes' at question '1 [D01]' (Are you currently an independent health researcher (in a position where you can apply for research grants as Principal Investigator) at a Canadian university or research institute?) *and* Answer was NOT '>7 years' at question '3 [D02]' (Excluding any eligible delays (e.g., parental or medical leave), how many years has it been since you were first appointed as an independent researcher?)

❗ Choose one of the following answers
Please choose **only one** of the following:

- ◯ None of these
- ◯ An adult with a development disability
- ◯ An older person (≥ 65 years)
- ◯ Not listed: ____________

#### Do you have significant community responsibilities in your current role? (e.g., serving as a Council Member for a First Nation band)

Only answer this question if the following conditions are met:
Answer was 'Yes' at question '1 [D01]' (Are you currently an independent health researcher (in a position where you can apply for research grants as Principal Investigator) at a Canadian university or research institute?) *and* Answer was NOT '>7 years' at question '3 [D02]' (Excluding any eligible delays (e.g., parental or medical leave), how many years has it been since you were first appointed as an independent researcher?)

❗ Choose one of the following answers
Please choose **only one** of the following:

- ◯ No
- ◯ Prefer not to answer
- ◯ Yes, provide details if desired:

[ ]

#### Are your contributions to the community recognized and considered in assessments of your performance?

Only answer this question if the following conditions are met:
Answer was 'Yes' at question '1 [D01]' (Are you currently an independent health researcher (in a position where you can apply for research grants as Principal Investigator) at a Canadian university or research institute?) *and* Answer was NOT '>7 years' at question '3 [D02]' (Excluding any eligible delays (e.g., parental or medical leave), how many years has it been since you were first appointed as an independent researcher?) *and* Answer was 'Other' at question '23 [D28]' (Do you have significant community responsibilities in your current role? (e.g., serving as a Council Member for a First Nation band))

❗ Choose one of the following answers
Please choose **only one** of the following:

- ◯ No, not at all
- ◯ Yes - partially
- ◯ Yes - fully

### Mental Health and Burnout

##### Oldenburg Burnout Inventory

#### The Oldenburg Burnout Inventory is a measure of burnout and has been used in previous studies of academic burnout.

#### All questions are answered on a scale of "Strongly Agree, Agree, Disagree, Strongly Disagree"

Only answer this question if the following conditions are met:
Answer was 'Yes' at question '1 [D01]' (Are you currently an independent health researcher (in a position where you can apply for research grants as Principal Investigator) at a Canadian university or research institute?) *and* Answer was NOT '>7 years' at question '3 [D02]' (Excluding any eligible delays (e.g., parental or medical leave), how many years has it been

since you were first appointed as an independent researcher?)

Please choose the appropriate response for each item:

| | Strongly Agree | Agree | Disagree | Strongly Disagree |
|---|---|---|---|---|
| **I always find new and interesting aspects in my work** | ○ | ○ | ○ | ○ |
| **There are days when I feel tired before I arrive at work** | ○ | ○ | ○ | ○ |
| **It happens more and more often that I talk about my work in a negative way** | ○ | ○ | ○ | ○ |
| **After work, I tend to need more time than in the past in order to relax** | ○ | ○ | ○ | ○ |
| **I can tolerate the pressure of work very well** | ○ | ○ | ○ | ○ |
| **Lately, I tend to think less at work and do my job almost mechanically** | ○ | ○ | ○ | ○ |
| **I find my work to be a positive challenge** | ○ | ○ | ○ | ○ |
| **During my work, I often feel emotionally drained** | ○ | ○ | ○ | ○ |
| **Over time, one can become disconnected** | ○ | ○ | ○ | ○ |

| | | | | |
|---|---|---|---|---|
| **from this type of work** | | | | |
| **After working, I have enough energy for my leisure activities** | ○ | ○ | ○ | ○ |
| **Sometimes, I feel sickened by my work tasks** | ○ | ○ | ○ | ○ |
| **After my work, I usually feel worn out and weary** | ○ | ○ | ○ | ○ |
| **This is the only type of work I can imagine myself doing** | ○ | ○ | ○ | ○ |
| **Usually, I can manage the amount of work well** | ○ | ○ | ○ | ○ |
| **I feel more and more engaged in my work** | ○ | ○ | ○ | ○ |
| **When I work, I usually feel energized** | ○ | ○ | ○ | ○ |

#### Has your mental health been impacted by the pandemic?

Only answer this question if the following conditions are met:
Answer was 'Yes' at question '1 [D01]' (Are you currently an independent health researcher (in a position where you can apply for research grants as Principal Investigator) at a Canadian university or research institute?) *and* Answer was NOT '>7 years' at question '3 [D02]' (Excluding any eligible delays (e.g., parental or medical leave), how many years has it been since you were first appointed as an independent researcher?)

❗ Choose one of the following answers
Please choose **only one** of the following:

- ◯ Yes, it has significantly worsened
- ◯ Yes, it has moderately worsened
- ◯ No, it has not changed
- ◯ Yes, it has improved moderately
- ◯ Yes, it has improved significantly
- ◯ Prefer not to answer

### Research and Grants

#### Did you alter your research direction to focus on COVID-19 and its impacts?

Only answer this question if the following conditions are met:
Answer was 'Yes' at question '1 [D01]' (Are you currently an independent health researcher (in a position where you can apply for research grants as Principal Investigator) at a Canadian university or research institute?) *and* Answer was NOT '>7 years' at question '3 [D02]' (Excluding any eligible delays (e.g., parental or medical leave), how many years has it been since you were first appointed as an independent researcher?)

❗ Choose one of the following answers
Please choose **only one** of the following:

- ◯ No
- ◯ Yes - Partially
- ◯ Yes - Fully

#### How many **tri-council** (i.e., SSHRC, CIHR, NSERC) grants did you apply for as principal applicant or investigator in the year before the COVID-19 pandemic (2019)?

Only answer this question if the following conditions are met:
Answer was 'Yes' at question '1 [D01]' (Are you currently an independent health researcher (in a position where you can apply for research grants as Principal Investigator) at a Canadian university or research institute?) *and* Answer was NOT '<1 year' *or* '6 years' *or* '5 years' *or* '4 years' *or* '>7 years' at question '3 [D02]' (Excluding any eligible delays (e.g., parental or medical leave), how many years has it been since you were first appointed as an independent researcher?)

❗ Only numbers may be entered in this field.
Please write your answer here:

#### How many **non-tri-council** grants did you apply for as principal applicant or principal investigator in the year before the COVID-19 pandemic (2019)?

Only answer this question if the following conditions are met:
Answer was 'Yes' at question '1 [D01]' (Are you currently an independent health researcher (in a position where you can apply for research grants as Principal Investigator) at a Canadian university or research institute?) *and* Answer was NOT '>7 years' *or* '4 years' *or* '5 years' *or* '6 years' *or* '<1 year' at question '3 [D02]' (Excluding any eligible delays (e.g., parental or medical leave), how many years has it been since you were first appointed as an independent researcher?)

❗ Only numbers may be entered in this field.
Please write your answer here:

#### How many successful grant applications did you have in 2019?

Only answer this question if the following conditions are met:
Answer was 'Yes' at question '1 [D01]' (Are you currently an independent health researcher (in a position where you can apply for research grants as Principal Investigator) at a Canadian university or research institute?) *and* Answer was NOT '4 years' *or* '5 years' *or* '6 years' *or* '<1 year' at question '3 [D02]' (Excluding any eligible delays (e.g., parental or medical leave), how many years has it been since you were first appointed as an independent researcher?)

❗ Only numbers may be entered in this field.
Please write your answer here:

#### How many **tri-council** (i.e., SSHRC, CIHR, NSERC) grants did you apply for as principal applicant or principal investigator in the first year of the pandemic (2020)?

Only answer this question if the following conditions are met:
Answer was 'Yes' at question '1 [D01]' (Are you currently an independent health researcher (in a position where you can apply for research grants as Principal Investigator) at a Canadian university or research institute?) *and* Answer was NOT '5 years' *or* '6 years' *or* '<1 year' at question '3 [D02]' (Excluding any eligible delays (e.g., parental or medical leave), how many years has it been since you were first appointed as an independent researcher?)

❗ Only numbers may be entered in this field.
Please write your answer here:

#### How many **non-tri-council** grants did you apply for as principal applicant or principal investigator in the first year of the pandemic (2020)?

Only answer this question if the following conditions are met:
Answer was 'Yes' at question '1 [D01]' (Are you currently an independent health researcher (in a position where you can apply for research grants as Principal Investigator) at a Canadian university or research institute?) *and* Answer was NOT '5 years' *or* '6 years' *or* '<1 year' at question '3 [D02]' (Excluding any eligible delays (e.g., parental or medical leave), how many years has it been since you were first appointed as an independent researcher?)

❗ Only numbers may be entered in this field.
Please write your answer here:

#### How many sucessful grant applications did you have in 2020?

Only answer this question if the following conditions are met:
Answer was 'Yes' at question '1 [D01]' (Are you currently an independent health researcher (in a position where you can apply for research grants as Principal Investigator) at a Canadian university or research institute?) *and* Answer was NOT '5 years' *or* '6 years' *or* '<1 year' at question '3 [D02]' (Excluding any eligible delays (e.g., parental or medical leave), how many years has it been since you were first appointed as an independent researcher?)

❗ Only numbers may be entered in this field.
Please write your answer here:

#### How many **tri-council** (i.e., SSHRC, CIHR, NSERC) grants did you apply for as principal applicant or principal investigator in the second year of the pandemic (2021)?

Only answer this question if the following conditions are met:
Answer was 'Yes' at question '1 [D01]' (Are you currently an independent health researcher (in a position where you can apply for research grants as Principal Investigator) at a Canadian university or research institute?) *and* Answer was NOT '>7 years' at question '3 [D02]' (Excluding any eligible delays (e.g., parental or medical leave), how many years has it been since you were first appointed as an independent researcher?)

❗ Only numbers may be entered in this field.
Please write your answer here:

#### How many **non-tri-council** grants did you apply for as principal applicant or principal investigator in the second year of the pandemic (2021)?

Only answer this question if the following conditions are met:
Answer was 'Yes' at question '1 [D01]' (Are you currently an independent health researcher (in a position where you can apply for research grants as Principal Investigator) at a Canadian university or research institute?) *and* Answer was NOT '>7 years' at question '3 [D02]' (Excluding any eligible delays (e.g., parental or medical leave), how many years has it been since you were first appointed as an independent researcher?)

❶ Only numbers may be entered in this field.
Please write your answer here:

#### How many successful grant applications did you have in 2021?

Only answer this question if the following conditions are met:
Answer was 'Yes' at question '1 [D01]' (Are you currently an independent health researcher (in a position where you can apply for research grants as Principal Investigator) at a Canadian university or research institute?) *and* Answer was NOT '6 years' *or* '<1 year' at question '3 [D02]' (Excluding any eligible delays (e.g., parental or medical leave), how many years has it been since you were first appointed as an independent researcher?)

❶ Only numbers may be entered in this field.
Please write your answer here:

#### How many **tri-council** (i.e., SSHRC, CIHR, NSERC) grants did you apply for as principal applicant or principal investigator in 2022?

Only answer this question if the following conditions are met:
Answer was 'Yes' at question '1 [D01]' (Are you currently an independent health researcher (in a position where you can apply for research grants as Principal Investigator) at a Canadian university or research institute?) *and* Answer was NOT '>7 years' at question '3 [D02]' (Excluding any eligible delays (e.g., parental or medical leave), how many years has it been since you were first appointed as an independent researcher?)

Only numbers may be entered in this field.
Please write your answer here:

#### How many **non-tri-council** grants did you apply for as principal applicant or principal investigator in 2022?

Only answer this question if the following conditions are met:
Answer was 'Yes' at question '1 [D01]' (Are you currently an independent health researcher (in a position where you can apply for research grants as Principal Investigator) at a Canadian university or research institute?) *and* Answer was NOT '>7 years' at question '3 [D02]' (Excluding any eligible delays (e.g., parental or medical leave), how many years has it been since you were first appointed as an independent researcher?)

Only numbers may be entered in this field.
Please write your answer here:

#### How many successful grant applications did you have in 2022?

❗ Only numbers may be entered in this field.
Please write your answer here:

#### Where have you submitted grant applications? SELECT ALL THAT APPLY

Only answer this question if the following conditions are met:
Answer was 'Yes' at question '1 [D01]' (Are you currently an independent health researcher (in a position where you can apply for research grants as Principal Investigator) at a Canadian university or research institute?) *and* Answer was NOT '>7 years' at question '3 [D02]' (Excluding any eligible delays (e.g., parental or medical leave), how many years has it been since you were first appointed as an independent researcher?)

❗ Check all that apply
Please choose **all** that apply:

- ☐ Tri-council agencies
- ☐ Other federal agencies (e.g., Genome Canada, New Frontiers in Research Fund)
- ☐ Non-profit/Charities (e.g., Canadian Cancer Society, Heart & Stroke Foundation)
- ☐ Provincial or territorial funding agencies
- ☐ Agencies based in the United States (e.g., National Institutes for Health)
- ☐ International agencies not based in the United States
- ☐ Not listed::

### If you have not applied for any grants as principal applicant or principal investigator or applied for fewer than you expected, why was that? (SELECT ALL THAT APPLY)

Only answer this question if the following conditions are met:
Answer was 'Yes' at question '1 [D01]' (Are you currently an independent health researcher (in a position where you can apply for research grants as Principal Investigator) at a Canadian university or research institute?) *and* Answer was NOT '>7 years' at question '3 [D02]' (Excluding any eligible delays (e.g., parental or medical leave), how many years has it been since you were first appointed as an independent researcher?)

❗ Check all that apply
Please choose **all** that apply:

- ☐ On leave for a significant portion of the past few years
- ☐ Already sufficiently funded
- ☐ Did not feel ready
- ☐ Felt discouraged
- ☐ Inadequate resources to support grant-writing (e.g., time, graduate students, institutional grant-writing supports)
- ☐ Supply chain issues impacted on setting up lab/research space
- ☐ I have applied for as many grants as principal applicant or principal investigator as expected
- ☐ Other: ____________

#### If you have not applied for any grants in a co-investigator or collaborator role or applied for fewer than you expected, why was that? (SELECT ALL THAT APPLY)

Only answer this question if the following conditions are met:
Answer was 'Yes' at question '1 [D01]' (Are you currently an independent health researcher (in a position where you can apply for research grants as Principal Investigator) at a Canadian university or research institute?) *and* Answer was NOT '>7 years' at question '3 [D02]' (Excluding any eligible delays (e.g., parental or medical leave), how many years has it been since you were first appointed as an independent researcher?)

❗ Check all that apply
Please choose **all** that apply:

- ☐ On leave for a significant proportion of the last few years
- ☐ Already sufficiently funded or already had sufficient fundings
- ☐ Did not feel ready
- ☐ Felt discouraged
- ☐ Inadequate resources to support grant-writing (e.g., time, graduate students, institutional grant-writing supports)
- ☐ Unable to obtain necessary data
- ☐ I have applied for as many grants as co-investigator or collaborator as I had expected to
- ☐ Other: ____________

### University-based initiatives

#### Did you employer offer any time extensions for tenure application in light of the pandemic?

Only answer this question if the following conditions are met:
Answer was 'Yes' at question '1 [D01]' (Are you currently an independent health researcher (in a position where you can apply for research grants as Principal Investigator) at a Canadian university or research institute?) *and* Answer was NOT '>7 years' at question '3 [D02]' (Excluding any eligible delays (e.g., parental or medical leave), how many years has it been since you were first appointed as an independent researcher?)

❗ Choose one of the following answers
Please choose **only one** of the following:

- ◯ No
- ◯ Yes
- ◯ My university does not have tenure track positions
- ◯ I don't know
- ◯ Prefer not to answer

#### What length of extension in the tenure application process did they offer in light of the pandemic?

Only answer this question if the following conditions are met:
Answer was 'Yes' at question '1 [D01]' (Are you currently an independent health researcher (in a position where you can apply for research grants as Principal Investigator) at a Canadian university or research institute?) *and* Answer was 'Yes' at question '43 [U01]' (Did you employer offer any time extensions for tenure application in light of the pandemic?)

❗ Choose one of the following answers
Please choose **only one** of the following:

- ◯ 1 year
- ◯ 2 years
- ◯ >2 years

#### Will you take advantage of the extension to the application for tenure timeline?

Only answer this question if the following conditions are met:
Answer was NOT '>7 years' at question '3 [D02]' (Excluding any eligible delays (e.g., parental or medical leave), how many years has it been since you were first appointed as an independent researcher?) *and* Answer was 'Yes' at question '43 [U01]' (Did you employer offer any time extensions for tenure application in light of the pandemic?)

❗ Choose one of the following answers
Please choose **only one** of the following:

- ◯ Definitely No (or Didn't)
- ◯ Probably No
- ◯ I haven't decided yet
- ◯ Probably Yes
- ◯ Definitely Yes (or Did)

#### Consider your work-life. How difficult are the following?

Only answer this question if the following conditions are met:
Answer was 'Yes' at question '1 [D01]' (Are you currently an independent health researcher (in a position where you can apply for research grants as Principal Investigator) at a Canadian university or research institute?) *and* Answer was '3 years' *or* '2 years' *or* '1 year' *or* '<1 year' at question '3 [D02]' (Excluding any eligible delays (e.g., parental or medical leave), how many years has it been since you were first appointed as an independent researcher?)

Please choose the appropriate response for each item:

| | **More difficult than expected** | **As difficult/easy as expected** | **Easier than expected** | **Not applicable** |
|---|---|---|---|---|
| **Communication with** | ◯ | ◯ | ◯ | ◯ |

| | | | | |
|---|---|---|---|---|
| **research collaborators** | | | | |
| **Communication with your university administration** | ○ | ○ | ○ | ○ |
| **Starting new studies/projects** | ○ | ○ | ○ | ○ |
| **Recruiting** | ○ | ○ | ○ | ○ |
| **Mentoring** | ○ | ○ | ○ | ○ |
| **Change in work routines** | ○ | ○ | ○ | ○ |
| **Accessing on-site resources** | ○ | ○ | ○ | ○ |
| **Ordering/receiving reagents/materials** | ○ | ○ | ○ | ○ |
| **Teaching** | ○ | ○ | ○ | ○ |
| **Emotional support for trainees** | ○ | ○ | ○ | ○ |

#### Consider your experience pre- and post-pandemic. How difficult are the following?

Only answer this question if the following conditions are met:
Answer was 'Yes' at question '1 [D01]' (Are you currently an independent health researcher (in a position where you can apply for research grants as Principal Investigator) at a Canadian university or research institute?) *and* Answer was '7 years' *or* '6 years' *or* '5 years' *or* '4 years' at question '3 [D02]' (Excluding any eligible delays (e.g., parental or medical leave), how many years has it been since you were first appointed as an independent researcher?)

Please choose the appropriate response for each item:

| | More difficult now | No different | Easier now | Not applicable |
|---|---|---|---|---|
| **Communication with research collaborators** | ◯ | ◯ | ◯ | ◯ |
| **Communication with your university administration** | ◯ | ◯ | ◯ | ◯ |
| **Starting new studies/projects** | ◯ | ◯ | ◯ | ◯ |
| **Recruiting** | ◯ | ◯ | ◯ | ◯ |
| **Mentoring** | ◯ | ◯ | ◯ | ◯ |
| **Change in work routines** | ◯ | ◯ | ◯ | ◯ |
| **Accessing on-site resources** | ◯ | ◯ | ◯ | ◯ |
| **Ordering/receiving reagents/materials** | ◯ | ◯ | ◯ | ◯ |
| **Teaching** | ◯ | ◯ | ◯ | ◯ |
| **Emotional support for trainees** | ◯ | ◯ | ◯ | ◯ |

#### How do you feel your citizenship status impacted on your experience as an early career health researcher in Canada during the COVID-19 pandemic? (e.g. visa delays, work permit renewals, COVID-related separation from family)

Only answer this question if the following conditions are met:
Answer was 'Yes' at question '1 [D01]' (Are you currently an independent health researcher (in a position where you can apply for research grants as Principal Investigator) at a Canadian university or research institute?) *and* Answer was NOT '>7 years' at question '3 [D02]' (Excluding any eligible delays (e.g., parental or medical leave), how many years has it been since you were first appointed as an independent researcher?)

❗ Choose one of the following answers
Please choose **only one** of the following:

- ◯ Impacted my experience very negatively
- ◯ Impacted my experience negatively
- ◯ Had no impact on my experience
- ◯ Impacted my experience positively
- ◯ Impacted my experience very positively

#### What best describes your current location of work?

Only answer this question if the following conditions are met:
Answer was 'Yes' at question '1 [D01]' (Are you currently an independent health researcher (in a position where you can apply for research grants as Principal Investigator) at a Canadian university or research institute?) *and* Answer was NOT '>7 years' at question '3 [D02]' (Excluding any eligible delays (e.g., parental or medical leave), how many years has it been since you were first appointed as an independent researcher?)

❗ Choose one of the following answers
Please choose **only one** of the following:

- ◯ Primarily work at the office
- ◯ Primarily work at home
- ◯ Work both at the office and at home
- ◯ Other ________

#### Is there anything else you'd like to share about your experience as an early career researcher in Canada during the COVID-19 pandemic?

Only answer this question if the following conditions are met:
Answer was 'Yes' at question '1 [D01]' (Are you currently an independent health researcher (in a position where you can apply for research grants as Principal Investigator) at a Canadian university or research institute?) *and* Answer was NOT '>7 years' at question '3 [D02]' (Excluding any eligible delays (e.g., parental or medical leave), how many years has it been since you were first appointed as an independent researcher?)

Please write your answer here:

We will be translating study findings to be of direct, practical use to university administrators, health research institutions, and decision-makers at health research funding agencies across Canada. Supporting Early Career Health Researchers to conduct research in health and protecting their ability to thrive in the Canadian research ecosystem: i) ensures that Canadians have access to context-specific, evidence-informed health care, ii) prevents attrition of highly-trained personnel via emigration, and iii) enhances the quality of education provided within Canada's post-secondary institutions.

Thank you for taking the time to complete our survey. If you are experiencing any distressing emotions, contact Wellness Together Canada (https://www.wellnesstogether.ca/en-CA?lang=en-ca)at 1-866-585-0445 or text WELLNESS to 741741. If you are not yet a member of the Association for Canadian Early Career Health Researchers (ACECHR), please consider joining here (http://www.acechr.ca/members.html) . The results of this study, including policy briefs and published manuscript(s), will be distributed to the ACECHR membership.

02.06.2023 – 10:21

Submit your survey.
Thank you for completing this survey.
